# INFLAMMATORY BOWEL DISEASE INDUCES α-SYNUCLEIN AGGREGATION IN GUT AND BRAIN

**DOI:** 10.1101/2022.01.26.477259

**Authors:** Ana M. Espinosa-Oliva, Rocío Ruiz, Manuel Sarmiento Soto, Antonio Boza-Serrano, Ana I. Rodriguez-Perez, María A. Roca-Ceballos, Juan García-Revilla, Marti Santiago, Sébastien Serres, Vasiliki Economopoulus, Ana E. Carvajal, María D. Vázquez-Carretero, Pablo García-Miranda, Oxana Klementieva, María J. Oliva-Martín, Tomas Deierborg, Eloy Rivas, Nicola R. Sibson, José L. Labandeira-García, Alberto Machado, María J. Peral, Antonio J. Herrera, José L. Venero, Rocío M. de Pablos

**Affiliations:** Instituto de Biomedicina de Sevilla (IBiS), Hospital Universitario Virgen del Rocío/CSIC/Universidad de Sevilla, Sevilla, Spain and Departamento de Bioquímica y Biología Molecular, Facultad de Farmacia, Universidad de Sevilla, Spain; Cancer Research UK and Medical Research Council Oxford Institute for Radiation Oncology, Department of Oncology, University of Oxford, Churchill Hospital, Oxford, UK; Experimental Neuroinflammation Laboratory, Department of Experimental Medical Science, Lund University, BMC B11, Lund 221 84, Sweden; Research Center for Molecular Medicine and Chronic Diseases (CIMUS), University of Santiago de Compostela, Health Research Institute (IDIS), 15782 Santiago de Compostela, Spain; Networking Research Center on Neurodegenerative Diseases (CIBERNED), Madrid, Spain; School of Life Sciences, University of Nottingham, Nottingham NG7 2UH, UK; Departamento de Fisiología, Facultad de Farmacia, Universidad de Sevilla, Spain; Departamento de Anatomía Patológica, Hospital Universitario Virgen del Rocío, Sevilla, Spain; Dementia Research Laboratory, Department of Experimental Medical Science, Lund University, BMC B11, Lund 221 84, Sweden

**Author notes:** Sharing senior authorship (José L. Venero, Rocío M. de Pablos). These authors contributed equally to this work (Rocío Ruiz, Manuel Sarmiento Soto, Antonio Boza-Serrano). Correspondingauthor: Ana M. Espinosa-Oliva. Postal address: Departamento de Bioquímica y Biología Molecular, Facultad de Farmacia, Universidad de Sevilla. CalleProfesorGarcía González, 2. 41012-Sevilla, Spain.

## Abstract

According to Braak’s hypothesis, it is plausible that Parkinsońs disease (PD) starts in the enteric nervous system (ENS) to spread the brain via the vagus nerve. Thus, we were wondering whether human inflammatory bowel diseases (IBD) can progress with appearance of pathogenic α-synuclein (α-syn) in the gastrointestinal tract and midbrain dopaminergic neurons. Analysis of human gastrointestinal tract sections from IBD patients demonstrated the presence of pathogenic phosphorylated α-syn in both myenteric (Auerbach’s) and submucosal (Meissner’s) plexuses. Remarkably, PD subjects exhibit α-syn pathology in identical gastrointestinal locations. Analysis of human midbrain sections from IBD subjects revealed a clear displacement of neuromelanin in some nigral neurons from the ventral mesencephalon, which were inherently associated with presence of α-syn aggregates reminiscent of pale bodies. We also used different dextran sodium sulfate (DSS)-based rat models of gut inflammation (subchronic and chronic) to study the appearance of phosphorylated α-syn inclusions in both Auerbach’s and Meissner’s plexuses (gut), and in dopaminergic neuritic processes (brain) along with degeneration of nigral dopaminergic neurons, which are considered classical hallmarks of PD. Vagotomized DSS-treated animals exhibited pathological α-syn in the gut but failed to show dopaminergic cells degeneration and α-syn aggregation in the ventral mesencephalon. Taken together, these results strongly suggest that Braak’s hypothesis is plausible.

## INTRODUCTION

Parkinson’s disease (PD) is an irreversible neurodegenerative disorder characterized by a selective and gradual degeneration of the dopaminergic neurons in the substantia nigra (SN) pars compacta ^1^. Lewy pathology in PD includes Lewy neurites and Lewy bodies (LB), whose main component is fibrillar α-synuclein (α-syn)^2, 3^. Braak and colleagues classified PD in different stages, based on the appearance of Lewy pathology^4^. The earliest appearance of pathological α-syn was found in the dorsal motor nucleus of the vagus nerve (DMNV) and in the anterior olfactory nucleus. Those observations prompted the authors to suggest that these brain areas might be starting points for brain pathology to reach the SN pars compacta^4, 5^. In a subsequent study, the authors hypothesized that PD pathogenesis can have an origin on the enteric nervous system (ENS),gaining access to the brain through the entire gastrointestinal(GI) tract, including axons of the myenteric (Auerbach’s) plexus and/or the submucosal (Meissner’s) plexus via postganglionic neurons ^6^. The prospect that pathogenic α-syn may spread from the gut to the brain to further cause degeneration of the nigrostriatal dopaminergic system is certainly attractive, and recent data from experimental animals support this view. Initial studies showed that injections of PD brain lysates containing different forms of α-syn into the intestinal wall of rats leads to retrograde transport via the vagal nerve to the DMNV^7^. Similarly, injections of either preformed fibrils of α-syn, or adenovirus expressing α-syn, in the enteric system of rodents and non-human primates induce spreading of α-syn pathology to the DMNV ^8, 9^. Moreover, in a recent study, injections of preformed fibrils of α-syn into the GI wall of rodents led to vagal transport of pathogenic α-syn to the DMNV, followed by further spreading to the brain and dopaminergic degeneration ^10^. These studies highlight the potential involvement of the gut-brain axis in spreading of α-syn pathology under human disease-like conditions, including PD.

Previous studies from our group have demonstrated that different forms of peripheral inflammation, especially that induced by gut inflammation, increases the death of nigral dopaminergic neurons ^11–13^. Intriguingly, recent cohort studies have shown an association between PD and inflammatory bowel diseases (IBDs)^14–18^. IBDs are common chronic intestinal diseases usually classified as ulcerative colitis (UC) and Crohn’s disease (CD). A typical feature of IBD is long-lasting systemic inflammation. Consequently, with all these precedents, we were wondering whether IBD is associated with the appearance of pathogenic α-syn in the GI tract and midbrain dopaminergic neurons (Supplementary Fig. 1). To this end, experimental analyses have been performed using human and animal models. Specifically, we have used the dextran sodium sulfate (DSS)-based rat model of UC to study the role of gut inflammation itself on dopaminergic degeneration in the SN and appearance of pathological α-syn in the gut-brain axis. Using this model, we found distinctive PD features in the SN including disruption of the blood-brain barrier (BBB), degeneration of nigral dopaminergic neurons and, strikingly, the appearance of pathological α-syn inclusions within tyrosine hydroxylase (TH) fibers and neurons. Vagotomy fully prevented the appearance of pathogenic α-syn in the ventral mesencephalon and the loss of dopaminergic neurons. Finally, to enhance the clinical impact of our study, we analyzed human sections from the GI system and the ventral midbrain of adult subjects previously diagnosed with IBD. We also provided evidence of the appearance of pathogenic insoluble α-syn aggregates in both the Auerbach’s and Meissner’s plexuses of subjects with UC; notedly, both plexuses also show α-syn aggregates in PD subjects ^19^. Importantly, we found the formation of thioflavin-S-labeled insoluble α-syn aggregates in highly depigmented dopaminergic neurons from subjects of both UC and CD. Our results revitalize the importance of the gut-brain axis as a pathway of pathological α-syn transmission.

## RESULTS

### Phosphorylated α-syn(P-α-syn) accumulates in the gut of rats after subchronic treatment with DSS

We first wanted to know if DSS-induced gut inflammation generates pathological α-syn aggregates in the colon. To this end, male rats underwent a 28 days subchronic DSS treatment. To determine whether P-α-syn accumulates in the gut of DSS-treated rats, we performed immunofluorescence assays on colon sections and found a significant increase in P-α-syn in the mucosa and muscular external layer of the colon in DSS-treated rats compared with controls (Fig. 1a, b, c; p <0.05). The co-localization of P-α-syn with the pan-neuronal marker ubiquitin C-terminal hydrolase (UCHL)-1 demonstrated that P-α-syn was present in the ENS, in both myenteric and submucosal plexuses of rats with DSS-induced colitis (Fig. 1a). Analysis revealed that P-α-syn was located in the mucosal nerve fibers around the crypts, and in the neuronal somas and nerve fibers of either the submucosa or the muscular external layer. P-α-syn staining was also observed in non-neuronal cells probably from the immune system or glial cells.

**Figure 1.**
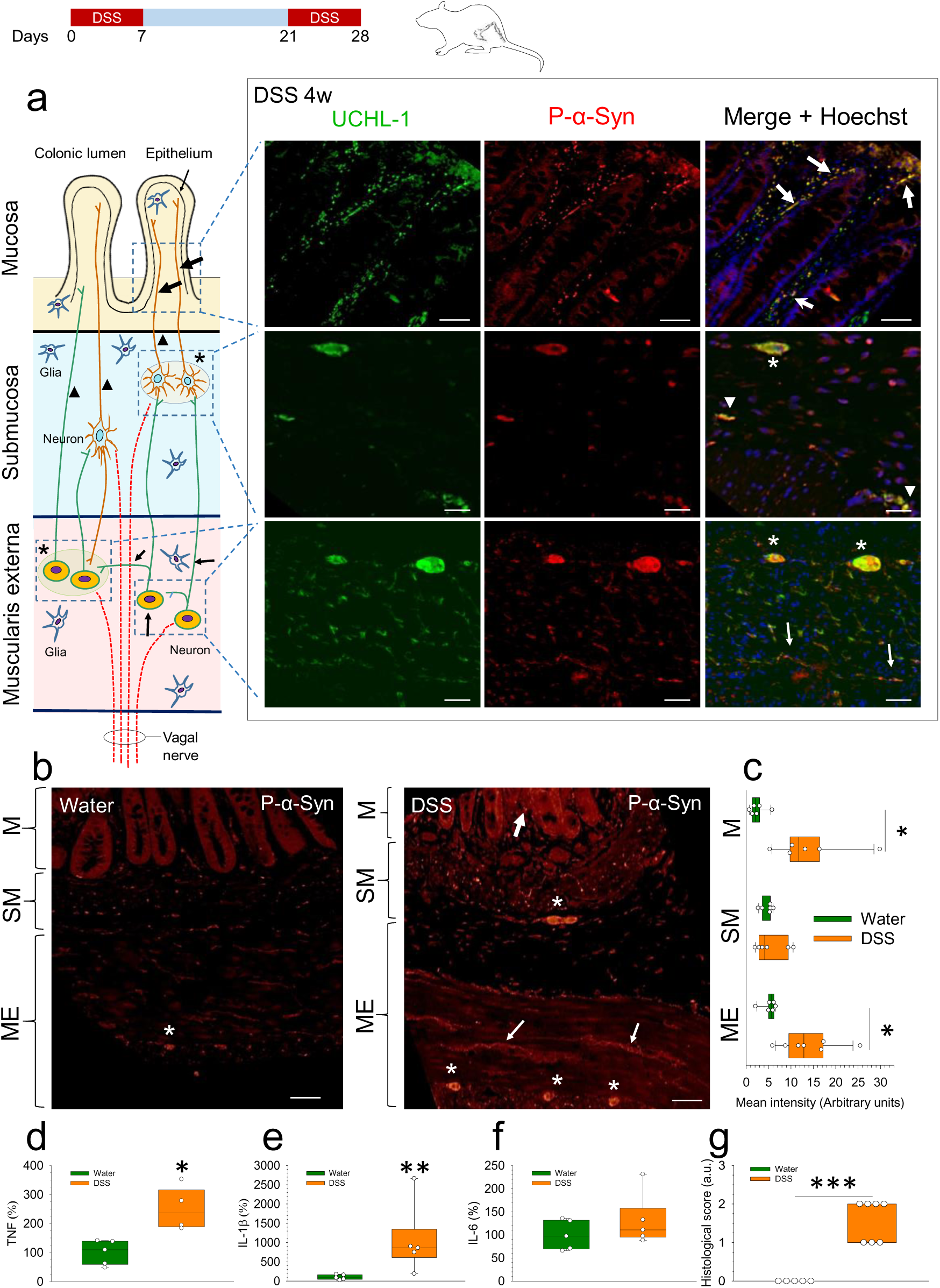
Immunolocalization of P-α-syn and upregulation of pro-inflammatory cytokines in the colon of rats under subchronic treatment with DSS. **(a)** Schematic cross-section of the colon wall illustrating the interconnected enteric plexuses, which contain neurons projecting to the mucosa, and vagal fibers. Co-localization of UCHL-1 and P-α-syn in the mucosal nerve fibers around a crypt (large arrows), in the submucosal nerve fibers (arrowheads) and ganglia (asterisks), and in the nerve fibers (small arrows) and ganglia (asterisks) of the muscular layer of a DSS-treated male rat. **(b)** Representative photographs of P-α-syn staining in the mucosa (M), submucosa (SM) and muscularis externa (ME) from colon of control (water) and DSS-treated male rats. Neuronal structures are indicated as in (a). Scale bars: 20 µm (a) and 50 µm (b). **(c)** Quantification of P-α-syn staining fluorescence intensity (as arbitrary units) measured in the mucosa (M), submucosa (SM) and muscularis externa (ME); N=6-7. **(d,e,f)** mRNA expression of TNF, IL1β and IL-6 quantified by RT-qPCR in the colon of control (water) and DSS-treated male rats; N=4-5. **(g)** Histological score of the colon of control (water) and DSS-treated male rats; N=5-7. Statistical analysis: Mann-Whitney U test for independent samples, with α=0.05; *, p< 0.05, **, p< 0.01 and ***, p < 0.001, comparing the DSS-treated with the control (water) group. Source data are provided as a Source Data file.

We also studied tumor necrosis factor (TNF), interleukin (IL)-1β and IL-6 gene expression in the gut using real-time quantitative reverse transcription PCR (RT-qPCR). In DSS-treated rats, there was an increase in TNF (252.6% of controls, p<0.05; Fig. 1d) and IL-1β (1073.8% of controls, p<0.01; Fig. 1e); IL-6 showed no significant changes (Fig. 1f). Histological damage of the colon of DSS-treated animals, assessed as a semiquantitative histological score (0 to 3, from healthy to severely damaged), showed a significant increase compared with control animals (Fig. 1g; p<0.001).

We also wanted to corroborate that DSS treatment actually induced peripheral inflammation. Hence, serum levels of TNF and IL-1β were measured using ELISA in animals receiving subchronic DSS treatment (Supplementary Fig. 2). This analysis showed that serum levels of TNF and IL-1β significantly increased in animals completing subchronic DSS treatment (Supplementary Fig.2a and b; p<0.05).

### Effects of the subchronic treatment with DSS on the brain

Using a stereological approach, we quantified TH-positive neurons in the SN of each animal after subchronic treatment with DSS. Intriguingly, analysis revealed that the right SN was systematically more affected than the left SN in animals treated with DSS, and hence stereological analysis was performed in the right and left SN. Notably, a significant reduced number (p<0.05) was evident in the right side (3924± 947 TH-positive cells) compared with the left side (6491 ± 731) of DSS-treated animals and with control animals (6847 ± 981 TH-positive cells in the left side; 6548 ± 619 in the right side; Fig. 2a, b and c). We also analyzed dopaminergic arborization by mean of immunofluorescence approach following subchronic DSS treatment and found a decrease in this parameter in the ventral mesencephalon of DSS-treated animals (Fig. 2e). These results indicate that the subchronic treatment with DSS causes unilateral damage in the right SN; this was confirmed in sections with TH-immunostaining and Nissl staining (Supplementary Fig. 3 a-e). In addition, we found that the intensity of a-syn staining in the TH-positive neurons was significantly greater in the midbrain of DSS-treated rats as compared with controls (363.5%; p<0.05; Fig. 2e), thus raising the possibility that α-syn aggregation in the brain is affected under conditions of gut inflammation. As a control, we measured the mean intensity of Hoechst in all sections, to detect a possible variation of tissue permeability between them and did not find any statistical difference (Fig. 2e). To confirm the presence of α-syn aggregates in the mesencephalon of rats subjected to DSS treatment, we took advantage of Fourier transform infrared (FTIR) spectroscopy, which allows us to measure abundance of β-sheet structures in the SN, a typical feature of amyloid deposits. FTIR spectra were recorded from left and right brain hemisphere sections. No significant differences in β-sheet structures between hemispheres were found in the control brain. However, levels of β-sheet structures were significantly elevated in the right SN from DSS-treated animals compared to the control group (p<0.01; Fig. 2f).

**Figure 2.**
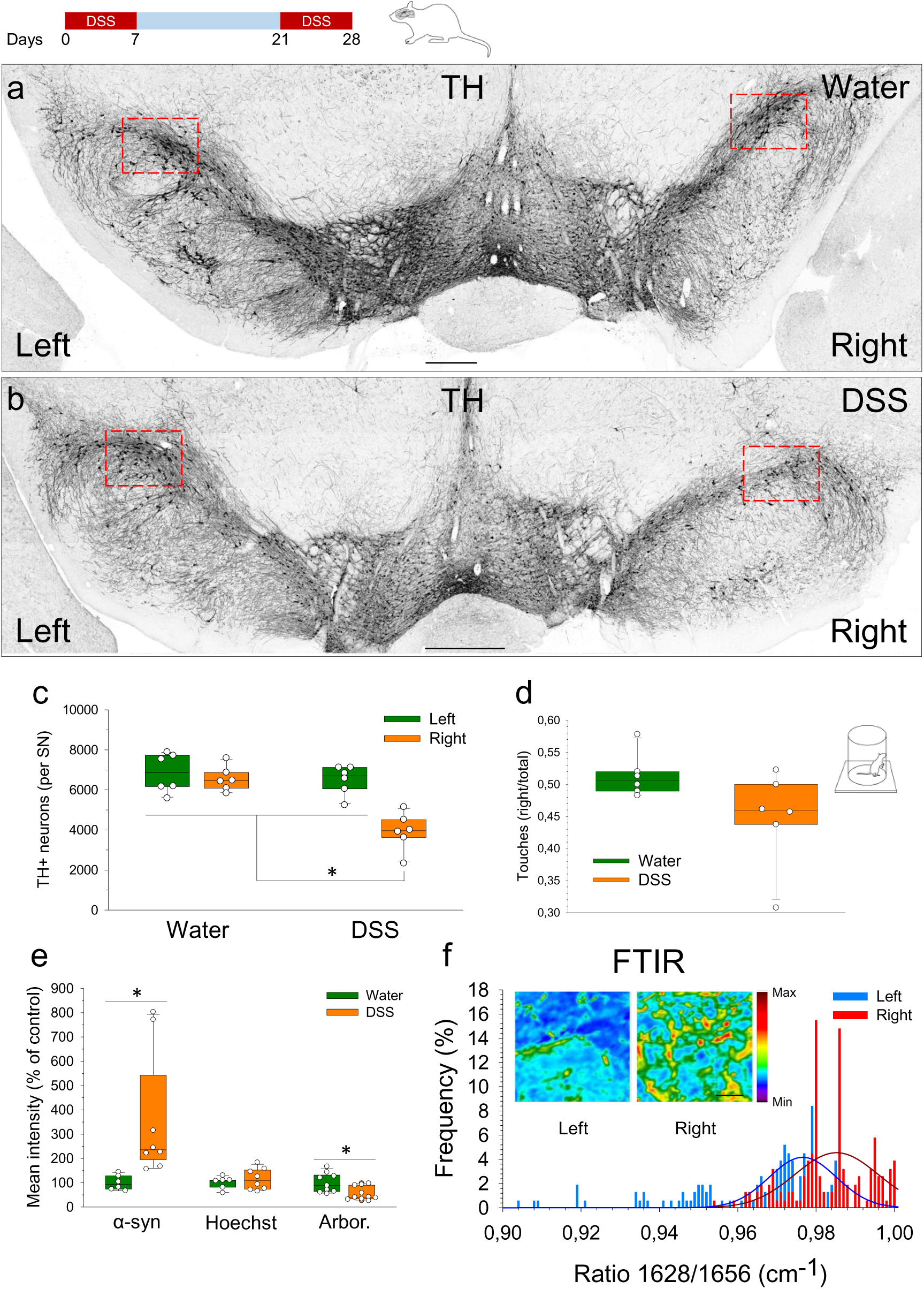
Effects of the subchronic treatment with DSS on the dopaminergic system, on dendritic arborisation and aggregation of a-syn in the rat brain of males. Coronal sections showing the whole SN from control **(a)** and animals treated with DSS **(b)**. TH immunohistochemistry illustrates a significant loss of dopaminergic cell bodies and dendrites in DSS-treated animals (see red boxes), which was particularly evident in the right SN. Scale bars: 1 mm. **(c)** Quantification of TH-positive cells in the SN of rats. Results are expressed as number of TH-positive neurons per SN. N=6 for all groups. Statistical analysis: The Kruskal-Wallis test for k independent measures followed by the Dunn’s post hoc test for pairwise comparisons with a Bonferroni correction with α=0.05: *, p<0.05 compared with the other groups. **(d)** Quantification of number of touches with the right and left paw (Cylinder test). N=6 for each group; results are expressed as ratio of touches (right/total) and were analyzed by the Mann-Whitney test; no significant difference was found. **(e)** Quantification of a-syn, Hoechst and arborization (Arbor.) intensity, using a section-by-section comparison between control and DSS-treated rats of the mean fluorescence intensity of these markers. Results are expressed as percentage of control values for each parameter; N=6-10. Statistical analysis: Mann-Whitney U test for two independent samples, with α=0.05. *, p< 0.05 compared with the control group. **(f)** Histogram showing the levels of β-sheet structures (recorded as FTIR spectra) in the left and right SN of rats treated with DSS for 28 days. Results are expressed as the ratio of β-sheet structures/total proteins (1628/1656 cm-1); N=155 for each side (5 animals, 33 measures per animal for each side). Statistical analysis: Mann-Whitney U-test for independent measures, with α=0.05. P value was 0.000 comparing the left with the right sides. Source data are provided as a Source Data file.

Our finding on TH-positive cells prompted us to further study motor behavior that could reflect asymmetric loss of nigral dopaminergic neurons. However, no statistical differences in DSS-treated animals compared to control animals were evident using the cylinder motor test (Fig. 2d). This result is in agreement with levels of dopamine (DA) and its metabolites, 3,4-dihydroxyphenylacetic (DOPAC) and homovanillic acid (HVA), measured in the striatum after the subchronic treatment with DSS, which also showed no statistical differences between control and DSS-treated animals (Supplementary Fig. 3f-h).

We also studied the expression of α-syn in other brain structures and found a qualitative increase in α-syn staining intensity in the DMNV on brain sections from DSS-treated animals in comparison to controls, with a greater increase evident on the right side (Supplementary Fig. 4). Other brain structures, such as the locus coeruleus and neocortex, were also studied, but no qualitative differences were evident (data not shown).

### BBB permeability and regional cerebral blood volume (rCBV) changes induced by DSS in rats

A single cohort of 6 rats underwent DSS treatment (1 week) and 7 consecutive magnetic resonance imaging (MRI) sessions to assess BBB integrity. Day 0 represented basal state of the BBB integrity and, thus, any MRI-based changes were compared to that time point thereafter. BBB permeability and rCBV changes were assessed daily by MRI in the nigrostriatal pathway of rats during the treatment (Fig. 3). Significant increases in rCBV within the SN were observed at day 3 (2.8-fold compared with day 0; Fig. 3e; p<0.01). No significant changes were found in the striatum (images not shown). However, an increase in mean signal intensity on T_1_-weighted images, an indicator of BBB disruption ^20^, was found in both the SN and striatum. In both regions, significant increases in BBB permeability were evident (see Fig. 3c, d and g for SN and Fig. 3 h for striatum) and peaked at day 4 (p<0.01). As explained above, in all cases mean signal intensity at day 0 was used as a baseline control. Finally, we aimed to evaluate whether BBB changes were different across hemispheres. Thus, MRI-based quantitative analysis was unilaterally performed and no significant differences in BBB permeability were found between in neither the left nor the right SN or in the striatum (Fig. 3i, j).

**Figure 3.**
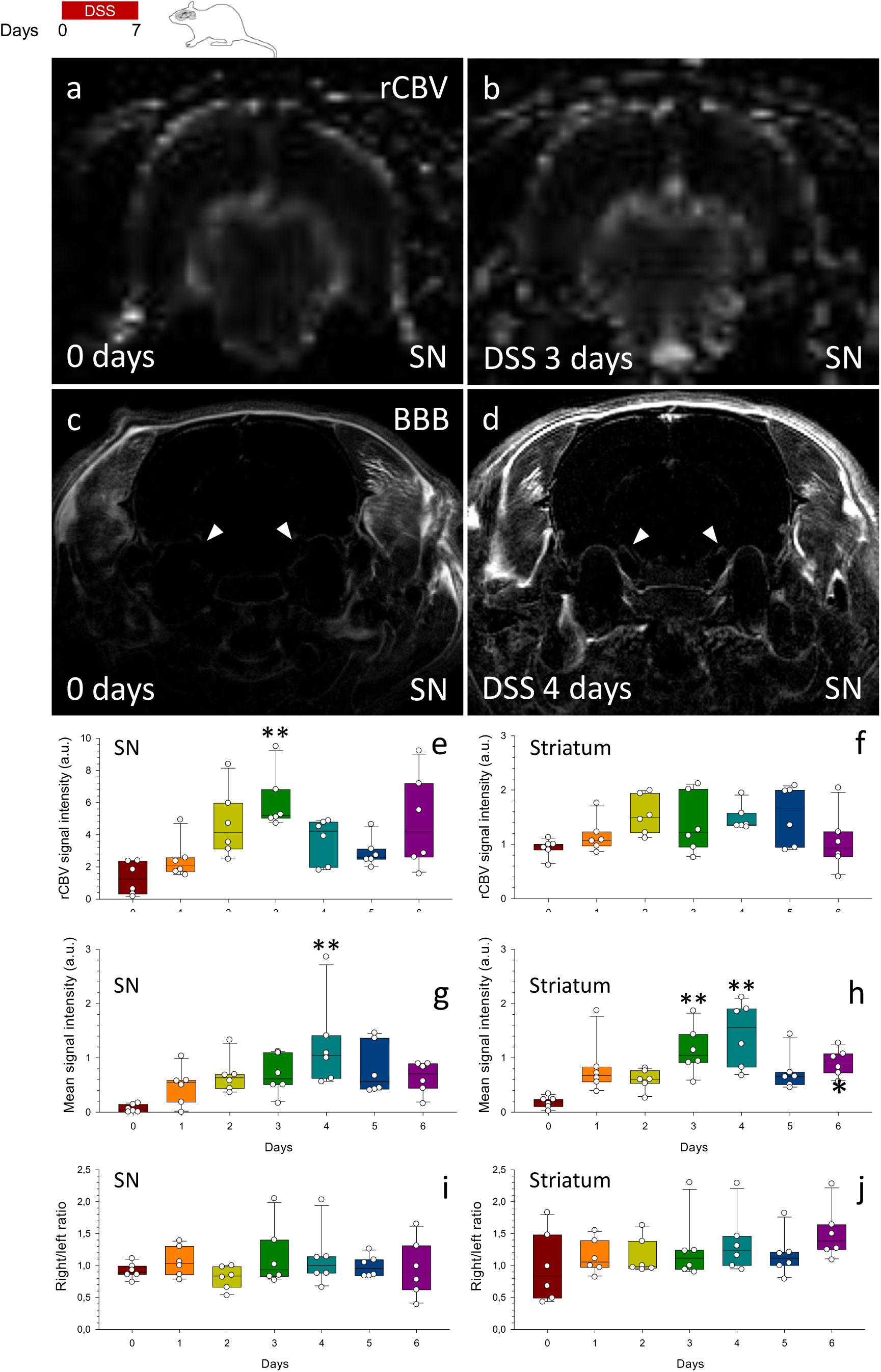
MRI study of SN and striatum of rats under DSS treatment. Representative rCBV maps showing the signal coming from the blood vessels within the SN area after 0 (**a**) and 3 days (**b**) of DSS treatment. Comparison of post-gadolinium T_1_-weighted images at day 0 (**c**) and day 4 (**d**) after DSS treatment show a clear enhancement within the SN 4 days after DSS treatment, suggesting changes in BBB structure and permeability. Arrowheads point SN. Quantification of rCBV within the SN (**e**) and the striatum (**f**) throughout the 6 days of DSS treatment. Quantification of the post-pre gadolinium-DTPA induced hyperintensity throughout DSS treatment within the SN (**g**) and striatum (**h**). (**i**) Measurement of gadolinium-DTPA enhancement comparing right versus left SN of DSS-treated rats. (**j**) As for (**i**), changes in right versus left Striatum. N=6. Statistical analysis: Friedman’s test for k related measures followed by post hoc test for pairwise comparisons with a Bonferroni correction with α=0.05. **, p< 0.01, compared with day 0. Source data are provided as a Source Data file.

### Peripheral immune cells infiltrate the brain of rats treated with acute DSS

Immune cells infiltration into brain parenchyma was analyzed by FACS to elucidate whether disruption of the BBB induced by DSS allows peripheral immune cells infiltration into the brain. To that aim, an intranigral injection of 2 µg of lipopolysaccharide (LPS) was used as positive control for detection of peripheral CD4^+^ and CD8^+^ lymphocytes, peripheral monocytes and macrophages and microglial cells in the SN (Supplementary Fig. 5).

Then, the effect of DSS treatment on peripheral cells infiltration into the SN was then assessed using FACS (Supplementary Fig. 6). Our results show that the number of T lymphocytes increased at day 2 (p<0.05), whilst the increase in monocytes infiltration was significantly altered at day 4 (p<0.05). Furthermore, microglial cells number was also increased at day 4 of DSS treatment (p<0.01). These data were further validated in a second cohort of animals, pooling 6 SN into each experimental group in order to increase the number of individuals and events. Again, T lymphocytes, monocytes and microglia within the SN were significantly increased after DSS treatments (day 2 and day 4 compared to control, Supplementary Fig. 6 d-f). Further analysis by immunofluorescence and dot blot demonstrated a significant increase in microglial activation (Iba1) (p<0.05), as well as numbers of CD4^+^ (p<0.05) and CD8^+^ (p<0.01) T lymphocytes within the SN three days after DSS treatment (Supplementary Fig. 6g-q).

### Microglial activation in rats under subchronic treatment with DSS

Once we demonstrated early changes in microglial population due to DSS, we evaluated microglial activation using Iba-1-positive cells in the SN at 3 days and after the subchronic DSS treatment (Supplementary Fig. 7). Notably, the number of microglia/macrophages increased significantly at day 3 (95.4% *vs.* controls, p<0.05; Supplementary Fig. 7g), and returned to control levels by 28 days after DSS treatment. Yet, around 30% cells retained the typical morphology of chronically activated microglia. Consequently, microglial morphology associated with different activation states was assessed in the SN in response to DSS treatment. This analysis showed a progression from quiescent microglia to amoeboid-like activate cells (Supplementary Fig. 8). After 3 days of DSS, 50% of the microglia/macrophages showed an activated morphology, with a reduction to 30% after 28 days. No differences were found between left and right SN.

Activated microglia release pro- or anti-inflammatory mediators ^21^, depending on their different phenotypes. Hence, we studied the activation of microglial cells at the molecular level by RT-qPCR for TNF, IL-1β, IL-6, arginase and IL-10 (Supplementary Fig. 7h-l). We also measured the expression of intercellular adhesion molecule (ICAM)-1 due to its role in cell infiltration (Supplementary Fig. 7m). Interestingly, we only found significant DSS-associated changes at week 4: TNF (221.3%, p<0.01), IL-1b (375.9%, p<0.01), IL-6 (750.4%, p<0.01), arginase-1 (228.2%, p<0.05), IL-10 (∼26-fold, p<0.01) and ICAM-1 (276.9%, p<0.01) compared with control values. This result supports the concept that systemic inflammation leads to fast and transient central changes^11, 22, 23^. The long-lasting inflammatory response in the ventral mesencephalon in response to DSS treatment may be indicative of neurodegenerative events.

### Vagotomy protects the dopaminergic system from the subchronic treatment with DSS and prevents midbrain inflammation

Since we found an increase in α-syn staining intensity in the DMNV (the entry point of the vagus nerve afferents to CNS) from DSS-treated animals, we next wanted to test the hypothesis that the gut-brain axis may be behind the loss of nigral TH neurons and α-syn aggregation under gut inflammatory conditions via the vagal nerve. To this end, we performed vagotomy on control and DSS-treated animals (Fig. 4). We found a clear protection of the dopaminergic system in vagotomized rats compared with sham-operated animals in terms of number of TH-positive neurons and arborisation (Fig.4c, f). We also found an increase (214.7%; p<0.05) in the intensity of α-syn in the TH-positive neurons in the midbrain from DSS-treated sham-operated animals that was abolished in DSS-treated vagotomized animals (Fig. 4d-f). Quantitative analysis in colon showed a significant increase of P-α-syn in the mucosa of sham-operated animals treated with DSS (422.2 %, p<0.05) that was prevented by vagotomy (Fig. 4g). In the muscular layer, P-α-syn increased in the DSS-treated animals, both sham-operated (402.8%, p<0.05) and vagotomized (469.4%, p<0.01), compared with the control animals (Fig. 4g). DSS treatment upregulated brain levels of inflammatory molecules such as TNF (667.8%, p<0.05), IL-1β (373.5%, p<0.01) and IL-6 (403.5%, p<0.01) as well as the anti-inflammatory cytokine IL-10 (235.9%, p<0.05) in DSS-treated animals compared with control animals (Fig. 4h-j, l), an effect prevented by vagotomy; arginase expression was not altered by DSS treatment (Fig. 4k). However, vagotomy did not prevent the upregulation of ICAM-1 (205.8%, p<0.01) by DSS (Fig. 4m). We conclude that gut inflammation induces degeneration of the nigral dopaminergic system with a significant major contribution of the vagal nerve. Indeed, vagotomy prevented death of dopaminergic neurons and upregulation of most pro-inflammatory markers in the ventral mesencephalon.

**Figure 4.**
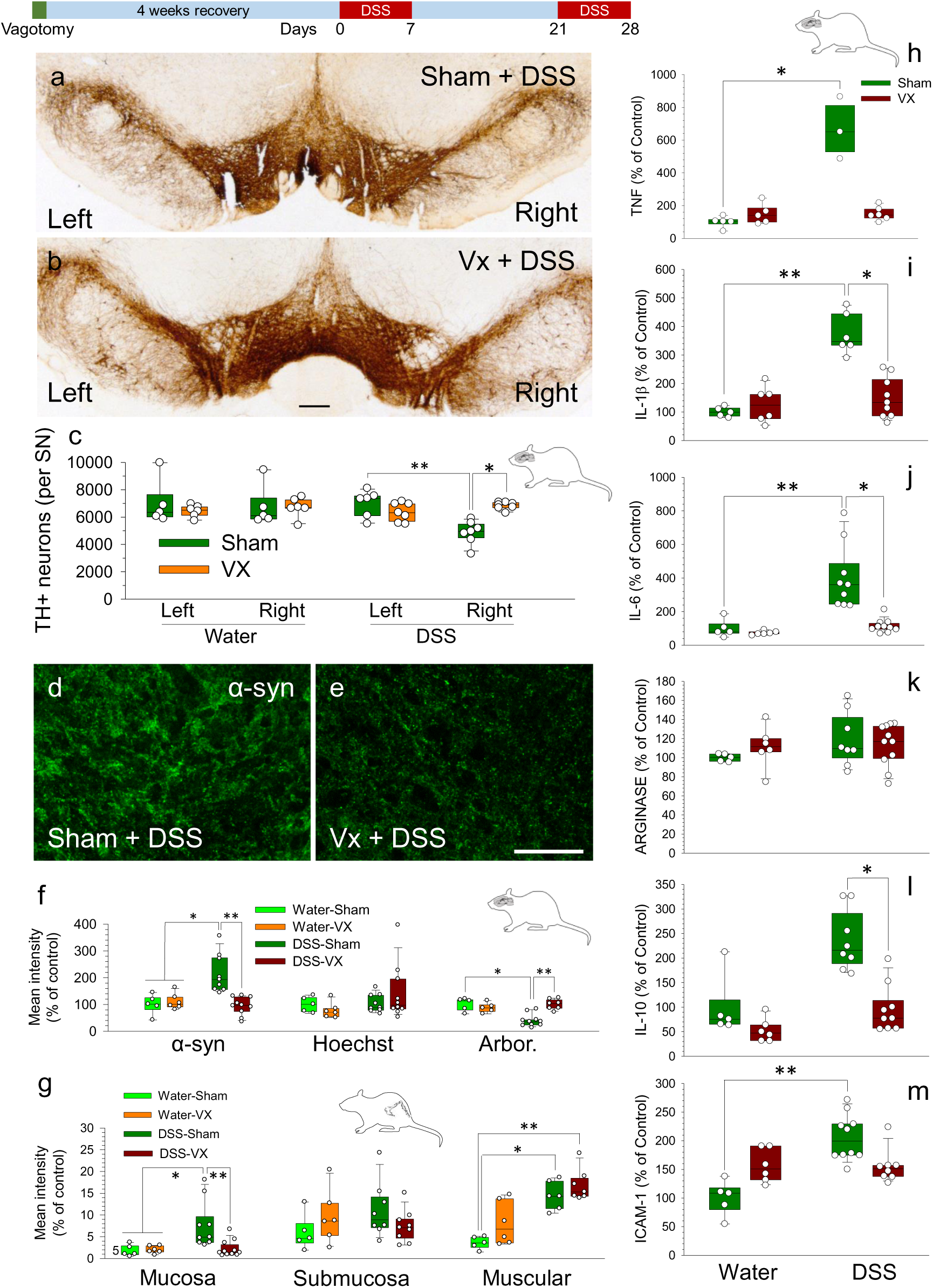
Effect of vagotomy on the changes induced in rats by the subchronic treatment with DSS. Coronal sections of TH immunohistochemistry showing the whole SN from (**a**) sham-operated and (**b**) vagotomized (Vx) animals, all of them treated with DSS. (**c**) Quantification of TH-positive cells in the SN of rats. Results are expressed as number of TH-positive neurons per SN; N=5-7. (**d**) Immunofluorescence shows an increase of a-syn in the SN of sham-operated animals treated with DSS that is reduced in the vagotomized rats (**e**). Scale bars: 1 mm (a and b) and 30 µm (d and e). (**f**) Quantification of the mean fluorescence intensity of P-a-syn, Hoechst and arborization (Arbor.) in the SN of rats. Results are expressed as percentage of control values for each parameter; N=5-10. (**g**) Quantification of fluorescence intensity of P-a-syn (as arbitrary units) measured in the mucosa, submucosa and muscularis externa (Muscular) in the colon of rats; N=5-11. Effect of DSS and vagotomy on the expression of TNF (**h**), IL-1β (**i**), IL-6 (**j**), arginase (**k**), IL-10 (**l**) and ICAM-1 (**m**) mRNAs. mRNA expression was quantified by RT-qPCR. Whilst DSS treatment increased the levels of these molecules in sham-operated animals, this effect was abolished when animals were vagotomized; N=3-11. Statistical analysis: Kruskal-Wallis test for k independent measures followed by the Dunn’s post hoc test for pairwise comparisons with a Bonferroni correction, with α=0.05: *, p<0.05 and **, p<0.01. Source data are provided as a Source Data file.

### Females do not show dopaminergic cell loss after the subchronic treatment with DSS

To study possible gender differences on the effect of DSS-induced colitis, we performed a new set of experiments with female rats (Supplementary Fig. 9). We found no statistical differences between control and DSS-treated female animals in terms of number of TH-positive neurons, motor test or intensity ofα-syn staining in the TH-positive neurons(Supplementary Fig. 9c-g).RT-qPCR analysis showed no differences in the expression levels of cytokines, arginase and ICAM-1 amongst any female experimental group in the SN (Supplementary Fig. 9 h-m).Quantitative analysis of P-α-syn showed a significant increase in the layers of the colon mucosa and muscularis externa, comparing DSS-treated and control female rats (Supplementary Fig. 10 a and c). Moreover, there were no differences in the markers of gut inflammation between male and female rats subjected to DSS treatment (Supplementary Fig. 10 d-g). All the experiments carried out to compare males with females were made with new cohorts of males and females that were treated at the same time.

### Effect of chronic treatment with DSS on the dopaminergic system, the aggregation of α–syn in brain and colon, and the inflammatory response in colon on rats

A typical feature of PD is asymmetry at the time of diagnosis ^24–26^ to further spread to the contralateral hemisphere ^27, 28^. Consequently, we wondered if the unilateral dopaminergic cell loss found in rats subjected to subchronic DSS treatment was an earlier event prior to acquiring bilateral cell loss. To this end, a chronic intermittent DSS treatment was implemented (3 weeks of DSS separated from each other by 2 weeks on water, plus an additional washout period of 2 weeks on water after the third week with DSS; see Supplementary Fig. 14b).

Using a stereological approach, we quantified TH-positive neurons in the right and left SN of each animal after the chronic treatment with DSS. Analysis revealed that both the left (3951 ± 493 TH-positive neurons) and the right SN (4114 ± 603) were affected (p<0.05 compared with both sides of controls Fig. 5d), contrary to the subchronic treatment with DSS that causes unilateral damage in the right SN.

**Figure 5.**
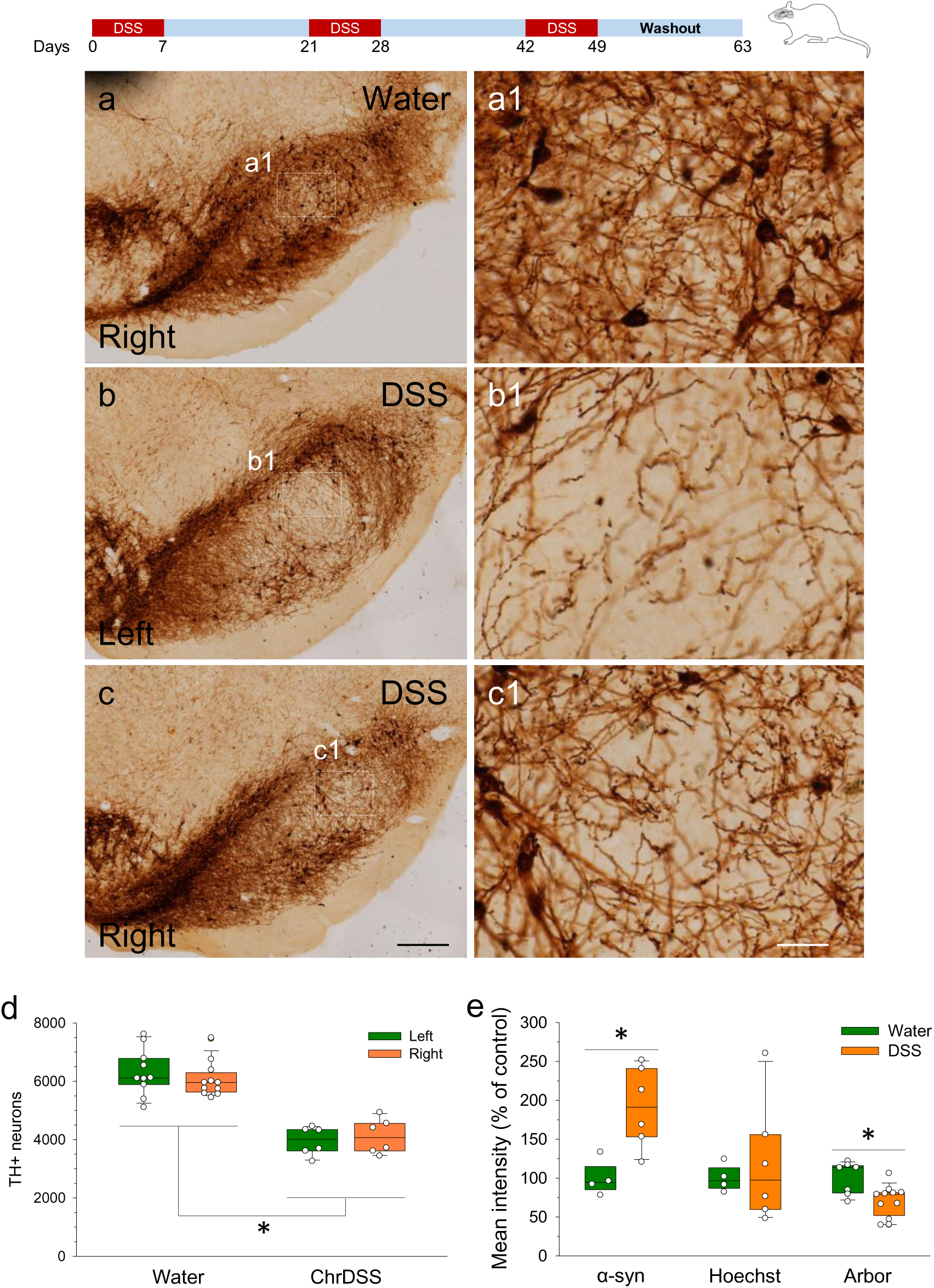
Effects of the chronic treatment with DSS on the dopaminergic system and dendritic arborisation in the rat brain. Coronal sections showing the whole SN from control animals (**a**) and animals treated with DSS (**b, c**). TH immunohistochemistry illustrates a significant loss of dopaminergic cell bodies and dendrites in DSS-treated animals (see white boxes and magnification in figures 5 a1-c1). Scale bars: 1 mm (a, b, c) and 25 µm (a1, b1, c1). (**d**) Quantification of TH-positive cells in the SN of rats under chronic treatments (ChrDSS) with DSS. Results are expressed as number of TH-positive neurons per SN. N=6 for all groups. Statistical analysis: Kruskal-Wallis test for k independent measures followed by the Dunn’s post hoc test for pairwise comparisons with a Bonferroni correction, with α=0.05: *, p<0.05. (**e**) Quantification of a-syn, Hoechst and arborization (Arbor.) intensity, using a section-by-section comparison of the mean fluorescence intensity of these markers between control rats and rats under chronic treatment with DSS. Results are expressed as percentage of control values for each parameter; N=6-10. Statistical analysis: Mann-Whitney U test for two independent samples, with α=0.05. *, p< 0.05 compared with the control group. Source data are provided as a Source Data file.

To confirm the presence of pathological α-syn in the ventral mesencephalon, we analyzed the presence of P-α-syn by confocal microscopy. In control animals, α-syn staining was diffuse corresponding to a classical presynaptic localization (Fig. 6 h-j). However, in DSS-treated animals, the presence of α-syn aggregates was mostly evident in dopaminergic processes (Fig. 6 k-m) but also in the soma (Fig. 6g). This analysis demonstrated the existence of P-α-syn dopaminergic neuritic inclusions in response to chronic DSS treatment (Fig. 6 n-p). In addition, we analyzed α-syn staining and dopaminergic arborization by mean of immunofluorescence approach regardless of the hemisphere following chronic DSS treatment (Figs. 5e). Overall, α-syn staining increased in the ventral mesencephalon of DSS-treated animals. We conclude that following DSS treatment, the right nigral dopaminergic system is preferentially affected at earlier stages and subsequently spreads to the contralateral one. This lateralization mimics the natural evolution from unilateral to bilateral damage in PD. In addition, our study revealed the existence of pathological α-syn neuritic inclusions, an event preceding the appearance of LB pathology ^29^.

**Figure 6.**
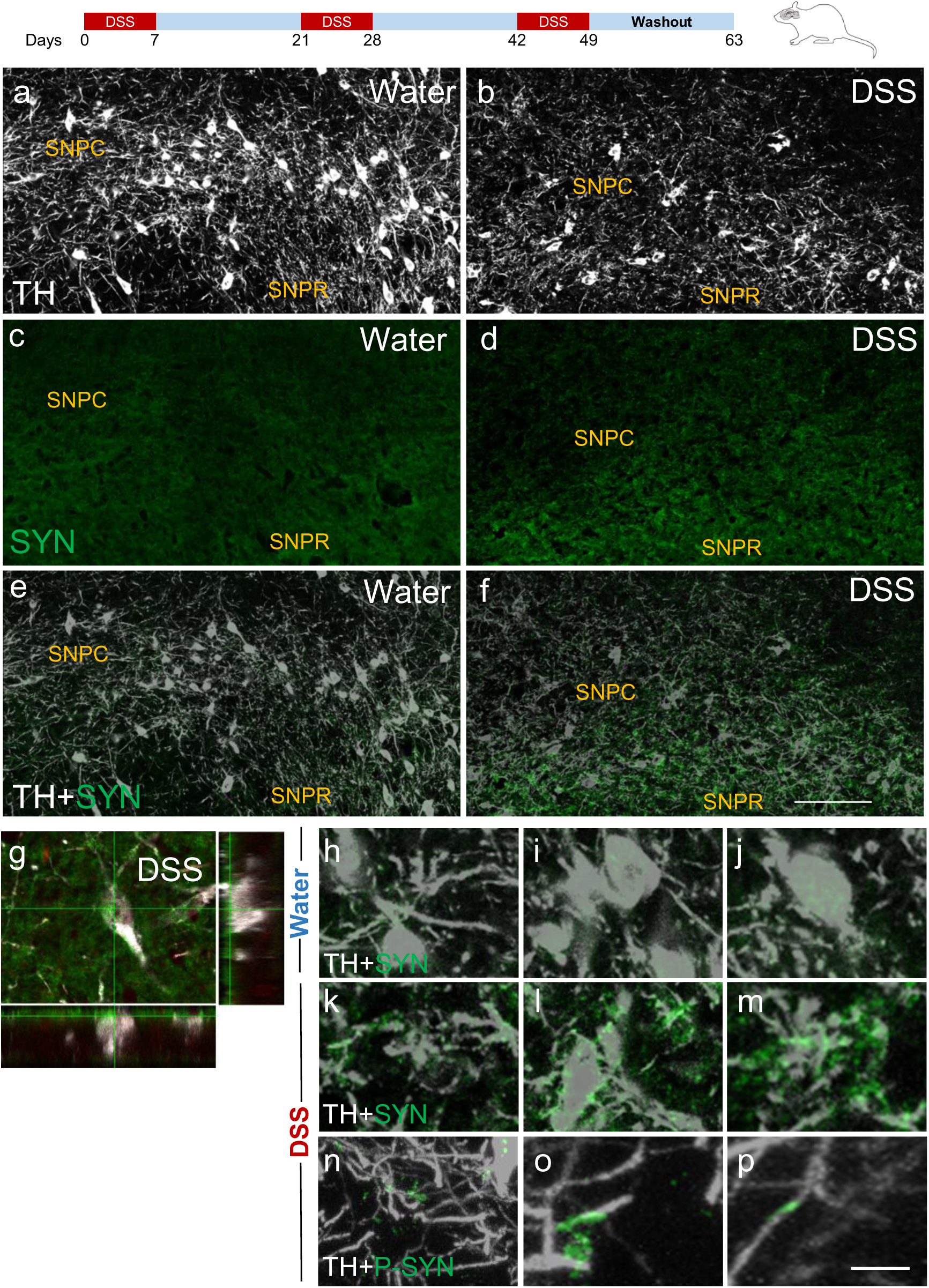
α-synuclein staining is highly increased in the SN from animals receiving chronic DSS treatment along with the appearance of aggregated forms. Coronal sections showing the SN.**a-e**: dual confocal immunohistochemistry for TH (**a** and**b**), α-syn (**c** and **d**) or both (**e** and **f**) from animals receiving either water (a,c,e) or DSS (b,d,f). **g-m**: higher-magnification photographs showing α-syn staining from both conditions. Note the absence of α-syn aggregates in control animals receiving water; the staining was clearly punctate in accordance with its presynaptic location (**h-j**). Aggregated forms of syn were evident in the SN of DSS-treated animals, especially within dopaminergic processes (**k-p**). Note the presence of α-syn aggregates in the soma of a dopaminergic neuron (g). The pathological nature of the α-syn inclusions was evidenced by P-α-syn staining (n-p). Scale bars: 200 µm (a-f) and 20µm (g-p).

In accordance to Braak’s hypothesis, Lewy pathology may start in enteric neurons of the gut ^10, 30, 31^. Our chronic DSS paradigm included a two-week washout period, an important feature to determine whether pathogenic α-syn aggregation and gut inflammation are long-lasting events. We found a significant increase in P-α-syn in the mucosa, submucosa and muscularis externa of the colon in DSS-treated rats compared with controls (Fig. 7a, b, c; p<0.05 for the three layers). This significant P-α-syn increase, compared with the subchronic treatment, in addition to the mucosa and muscularis externa, was also observed in the submucosa. The P-α-syn localization in the colon layers was similar to the subchronic treatment, in both plexuses of the ENS as showed by the co-localization with UCHL-1 and exhibited the same distribution in the neuronal soma and fibers (Fig. 7a, b). These results demonstrate that the DSS chronic treatment also causes α-syn pathology in the GI tract.

**Figure 7.**
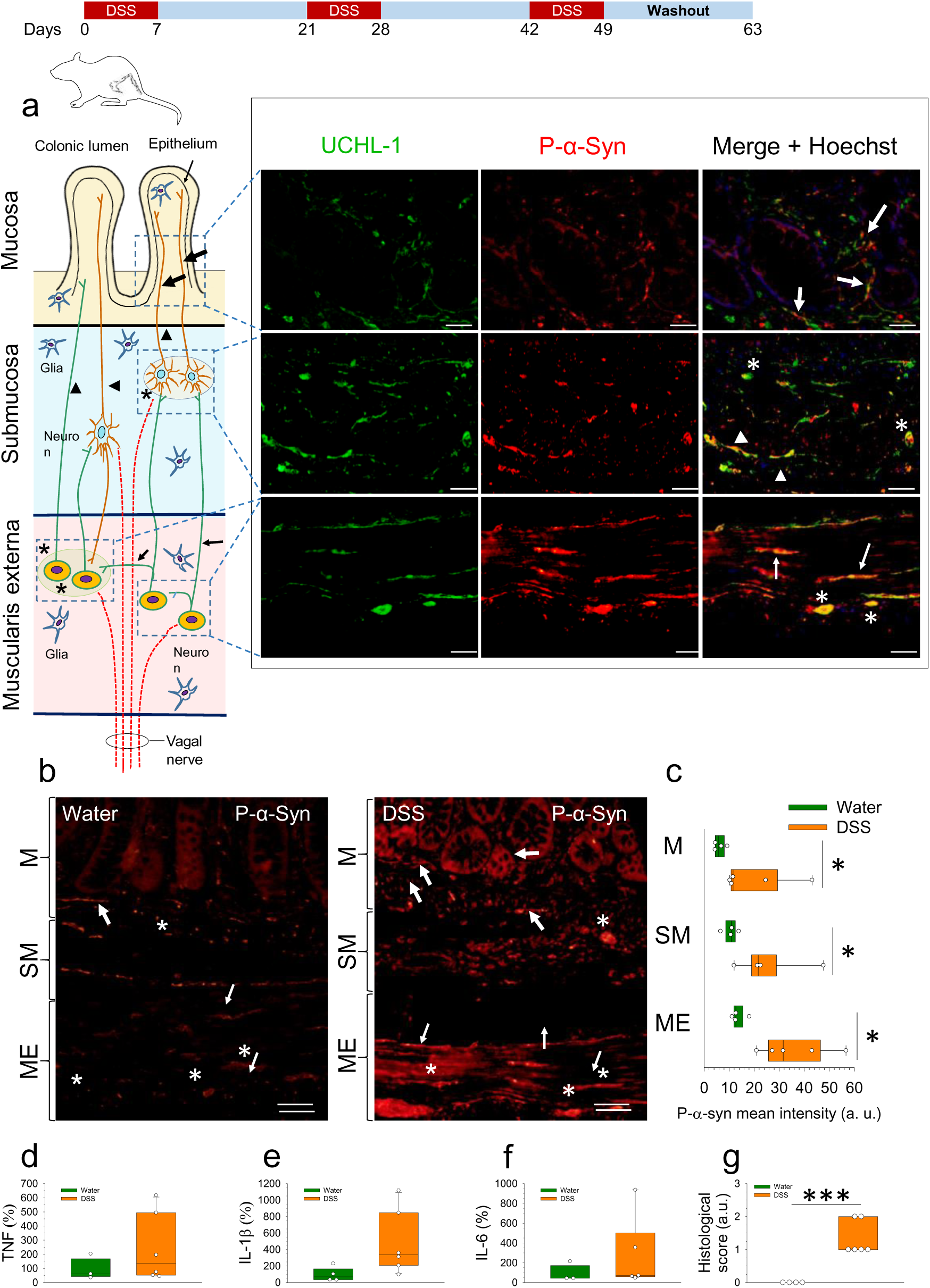
Immunolocalization of P-α-syn and upregulation of pro-inflammatory cytokines in the colon of rats under chronic treatment with DSS. **(a)** Schematic cross-section of the colon wall illustrating the interconnected enteric plexuses, which contain neurons projecting to the mucosa, and vagal fibers. Co-localization of UCHL-1 and P-α-syn in the mucosal nerve fibers around a crypt (large arrows), in the submucosal nerve fibers (arrowheads) and ganglia (asterisks), and in the nerve fibers (small arrows) and ganglia (asterisks) of the muscular layer of a DSS-treated male rat. **(b)** Representative photographs of P-α-syn staining in the mucosa (M), submucosa (SM) and muscularis externa (ME) from colon of control (water) and DSS-treated male rats. Neuronal structures are indicated as in (a). Scale bars: 20 µm (a) and 50 µm (b). **(c)** Quantification of fluorescence intensity (as arbitrary units) measured in the mucosa (M), submucosa (SM) and muscularis externa (ME); N=3-6. (**d, e, f**) mRNA expression of TNF, IL1β and IL-6 quantified by RT-qPCR in the colon of control (water) and DSS-treated male rats; N=4-5. (**g**) Histological score of the colon of control (water) and DSS-treated male rats; N=4-6. Statistical analysis: Mann-Whitney U test for independent samples, with α=0.05; *, p< 0.05 and ***, p < 0.001, comparing the DSS-treated with the control (water) group. Source data are provided as a Source Data file.

Regarding gut inflammation, we studied by RT-qPCR the levels of TNF, IL-1β and IL-6 mRNAs. Contrary to the subchronic treatment, no statistical difference was found between control and chronic DSS treatment (Fig. 7d-f) in accordance with the two-week washout period associated to the DSS paradigm. However, histological damage of the colon of DSS-treated animals showed a significant increase compared with the control animals (Fig. 7g; p<0.001).

ELISA analysis showed that, contrary to the subchronic treatment, serum levels of IL-1β were close to control values in animals receiving chronic DSS treatment, a feature likely related to the two-week washout period (Supplementary Fig. 2a). However, serum levels of TNF were clearly induced by the chronic DSS treatment (Supplementary Fig. 2b; p<0.05). This is of special interest since TNF seems to be critical for the transfer of inflammation from the periphery to the CNS ^32^ and highlight the long-lasting effect of gut inflammation into the periphery.

### Detection of P-α-syn in the gut of human subjects with UC

To investigate whether IBD could trigger α-syn aggregation in the ENS, we collected colon samples from eight IBB subjects and eight aged-matched controls (see Supplementary Table 1). This cohort included young (<50 years old) and old (>50 years old) subjects. We analyzed P-α-syn accumulation in the inflamed colon (Fig. 8) and found an increase in UC subjects compared to controls. Quantitative analysis of P-α-syn showed a significant increase in all layers of the colon wall (mucosa, submucosa and muscularis externa), compared with control subjects. No significant differences were evident between young (Fig. 8) and elderly UC patients (Supplementary Fig. 11).

**Figure 8.**
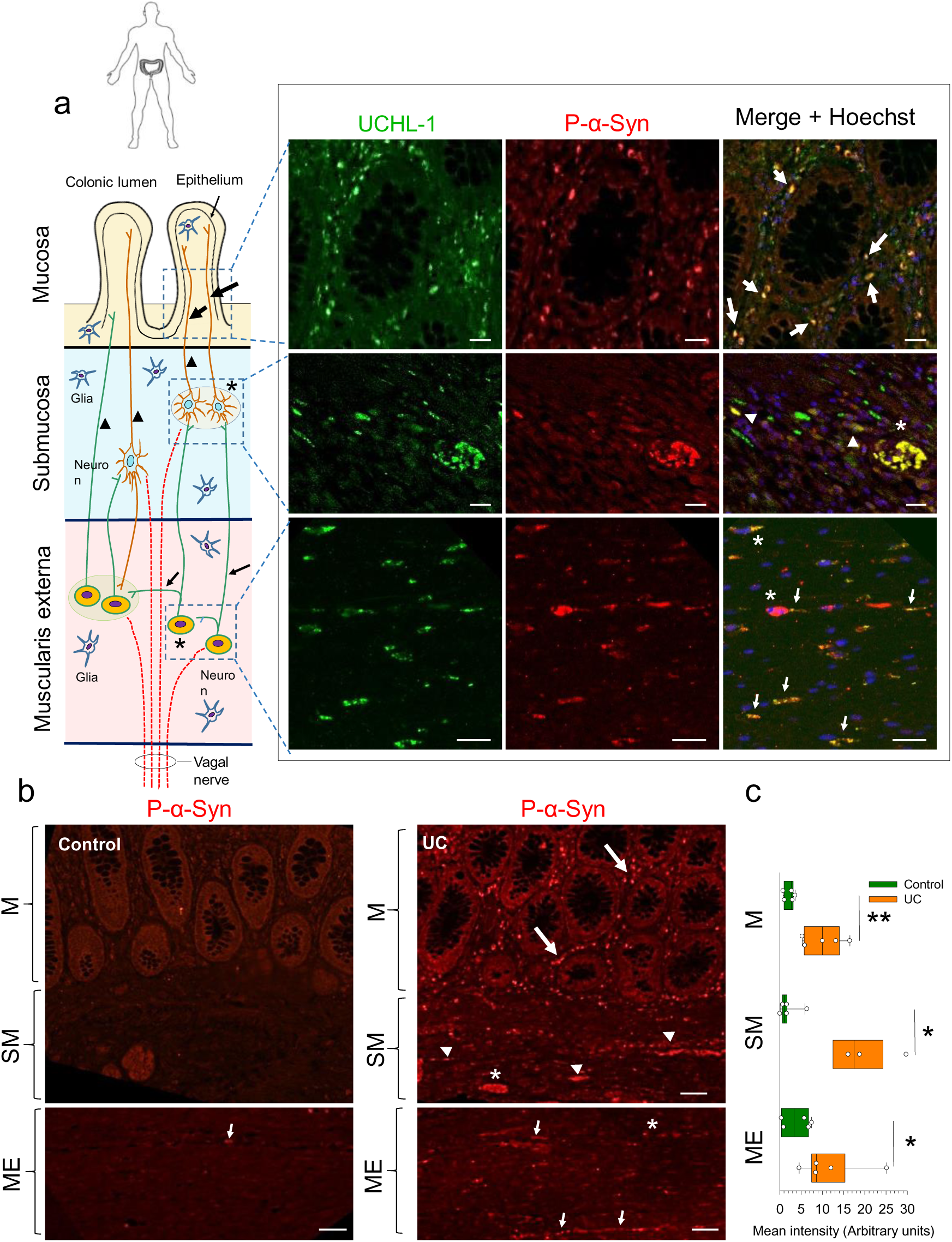
Immunolocalization of P-α-syn in the colon of human subjects with UC. (**a**) Schematic cross-section of the colon wall illustrating the interconnected enteric plexuses, which contain neurons projecting to the mucosa, and vagal fibers. Co-localization of P-α-syn and UCHL-1 in the mucosal nerve fibers around a crypt (large arrows), in the submucosal nerve fibers (arrowheads) and ganglia (asterisks), and in the neuronal somas (asterisks) and nerve fibers (small arrows) of the muscular layer of a 36-year-old patient with UC. P-α-syn is located in the mucosal nerve fibers around the crypts, in the submucosal neuronal somas and nerve fibers, and in the neuronal somas and nerve fibers of the muscular layer. Note that P-α-syn staining was also observed in non-neuronal cells. (**b**) Representative photographs of P-α-syn staining in the mucosa (M), submucosa (SM) and muscularis externa (ME) from colon of a control patient and a patient with UC. Neuronal structures are indicated as in (a). Scale bars: 20 µm (a) and 50 µm (b). (**c**) Quantification of fluorescence intensity (as arbitrary units) measured in the mucosa (M), submucosa (SM) and muscularis externa (ME); N=4-6. Statistical analysis: Mann-Whitney U test for independent samples, with α=0.05; *, p< 0.05 and **, p< 0.01 comparing the UC with the control group. Source data are provided as a Source Data file.

The co-localization of P-α-syn with UCHL-1 showed that P-α-syn was present in the ENS, in both myenteric and submucosal plexuses of subjects with UC. Our results demonstrated that P-α-syn is located in the mucosal nerve fibers around the crypts, in the submucosal neuronal somas and nerve fibers, and in the neuronal somas and nerve fibers of the muscular layer (Fig. 8a). We conclude that the colon of UC patients presents a P-α-syn expression and localization pattern similar to that of rats subjected to DSS subchronic or chronic treatment. Remarkably, this precise distribution pattern of α-syn pathology in the GI tract has been observed in PD patients thus supporting the involvement of the gut brain axis in spreading Lewy body pathology in the brain.

### Pathological forms of a-syn aggregate in the ventral mesencephalon of human IBD brains

Having established the existence of pathogenic α-syn in the inflamed gut of UC patients, we next wanted to determine whether a transition from healthy to altered neurons was evident in IBD subjects. In the human samples, dopaminergic neurons were first visualized by neuromelanin pigmentation. Early stages of α-syn pathology within nigral dopaminergic neurons include neuromelanin-related depigmentation and appearance of pale bodies. Remarkably, whilst neurons in the control subjects showed a normal pattern of neuromelanin pigmentation, we found clear displacement of neuromelanin in some neurons from the ventral mesencephalon in the eight IBD subjects, which were inherently associated to the presence of α-syn aggregates reminiscent of pale bodies (Fig. 9 and Supplementary Fig. 12). The pathological nature of the aggregates was demonstrated by the presence of phosphorylated S129 α-syn, the form of α-syn most clearly associated with synucleinopathies in both humans and animal models of PD (Fig. 9 and Supplementary Fig. 12; for review, see ^33^). Neurons from control subjects showed a normal pattern of α-syn, without inclusions. We have also quantified the expression of α-syn and P-α-syn in dopaminergic neurons and found an increase of 7.5 and 3.4-fold, respectively, compared to control subjects (p<0.05; Supplementary Fig. 12 m, n).

**Figure 9.**
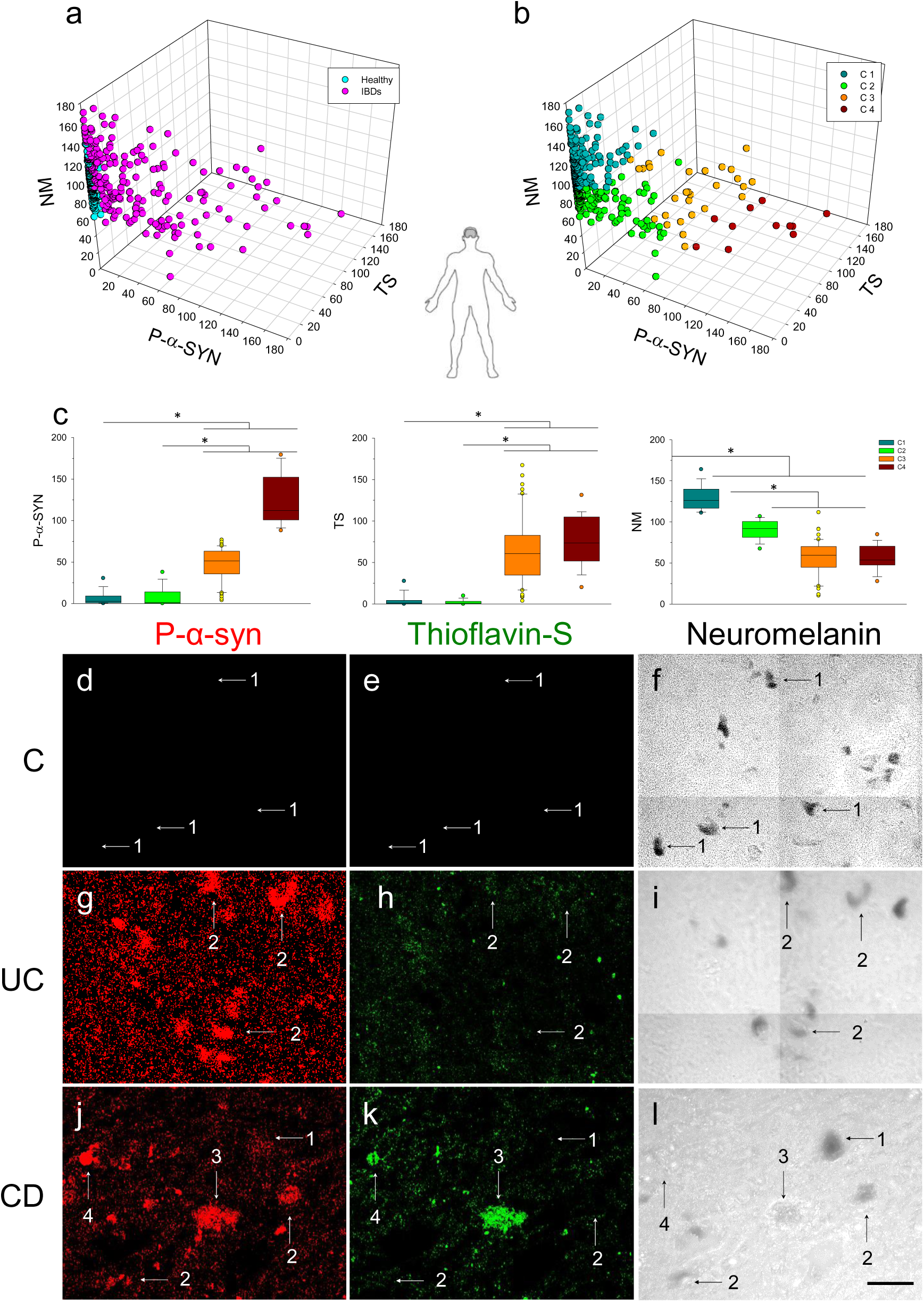
Analysis of mesencephalic neurons of healthy individuals and affected by IBDs. (**a**) Three-dimensional representation of the neurons according to their values of P-α-syn, thioflavin-S (TS) and neuromelanin (NM) expressed as mean intensity. The neurons from healthy individuals are shown in cyan and those from individuals with IBDs in pink. (**b**) Three-dimensional representation of the neurons grouped in four clusters after a hierarchical cluster analysis following a k-means method. (**c**) Boxes and whiskers representation of the three parameters used. Statistical analysis: Kruskal-Wallis test for k independent measures followed by the Mann-Whitney post hoc U test for pairwise comparisons with a Bonferroni correction, with α=0.05. *, p< 0.05. (**d-l**) Photomicrographs from the SN of healthy control (**d-f**) and IBD subjects (**g-l**) showing neurons belonging to each of the four clusters. Left panels show immunofluorescence to P-α-syn (red), central panels show immunofluorescence to thioflavin-S (green) and right panels show bright fields with neuromelanin as dark structures. The numbers denote clusters 1 to 4. Scale bar: 50 µm. Abbreviations: C, control; UC, ulcerative colitis; CD, Crohn’s disease. Source data are provided as a Source Data file.

Since pathogenic a-syn includes insoluble aggregates, we measured the levels of P-a-syn, thioflavin-S (insoluble aggregates) and neuromelanin in mesencephalic neurons from postmortem samples of healthy individuals and IBD patients (Fig. 9). The hierarchical cluster analysis of these neurons, following a k-means method, suggested that they clustered in 4 different groups showing, from cluster 1 to 4, an increase of P-a-syn and thioflavin-S and a decrease of neuromelanin. The clusters represent a gradient from healthy neurons characterized by low values of P-a-syn and thioflavin-S and high values of neuromelanin to neurons characterized by high values of P-a-syn and thioflavin-S, and low values of neuromelanin, which could suggest altered functionality. Neurons from healthy individuals are found only in clusters 1 and 2, whereas neurons from IBDs patients are found in all the clusters, indicating a wide range of neuronal functional states from healthy to a damaged state in these patients. Finally, to further study the nature of the observed pathologies, we performed a double immunostaining with P-α-syn and ubiquitin on human brain sections to qualitative evaluate if there was any difference (Supplementary Fig. 13). Notably, we found an increase in ubiquitin staining in IBD patients compared with controls. Furthermore, clear co-localization was evident between P-a-syn and ubiquitin, two Lewy pathology markers. Overall, our study demonstrates that IBD brains exhibit early features typically associated to a-syn pathology including neuronal depigmentation, neuromelanin displacement and appearance of aggregates reminiscent of pale bodies.

## DISCUSSION

It has been long established that most PD patients in all stages of the disorder exhibits α-syn pathology in the GI tract^34, 35^. Recent reports have established well-defined features of PD outside the brain, particularly in the GI tract including enteric neuropathology, GI dysfunction and mild inflammation^36^. Consequently, the possibility that PD starts in the ENS and then spreads to the brain via the vagus nerve is consistent with Braak’s hypothesis (bottom-up pathology). In this study, we have analyzed the GI tract and ventral midbrain from subjects previously diagnosed with IBD in terms of α-syn pathology. Our data sustains the existence of pathogenic α-syn in both gut and brain, thus, supporting the potential role of the ENS as a contributing factor in PD etiology. In addition, we have analyzed the effect of different DSS-based rat models of gut inflammation to demonstrate the appearance of P-α-syn inclusions in both Auerbach’s and Meissner’s plexuses (gut), ii) the appearance of P-α-syn inclusions in dopaminergic neuritic processes (brain) and iii) the degeneration of nigral dopaminergic neurons, which all are considered classical hallmarks in PD. In addition, in our experimental models, female rats showed neither neurodegeneration, nor α-syn increase or aggregation in the SN, in response to DSS treatment. Indeed, in the early stages of the disease, women usually shown a more benign PD phenotype than men, probably because of the neuroprotective role of estrogens^37^. Yet, analysis of human GI tract sections from both sexes, obtained from adult subjects diagnosed with UC, showed P-α-syn inclusions in both Auerbach’s and Meissner’s plexuses.

The concept of trans-synaptic α-syn spreading has been intensively investigated since the discovery that grafted embryonic dopaminergic neurons develop LB long after being transplanted in the striatum of PD subjects^38–40^. Different studies have consistently demonstrated that different α-syn fibril conformers may spread trans-synaptically and cause seeding of α-syn in the receiving neuron ^41, 42^. This concept has been well documented after injecting different types of α-syn aggregates into different brain areas, particularly the striatum ^41, 42^. However, convincing evidence supporting of α-syn spread pathology from the gut to the brain has remained elusive until recently ^7–10^. Previous reports have demonstrated that different forms of α-syn, including PD brain lysates, monomers, oligomers, preformed fibrils and adenoviruses overexpressing α-syn, are retrogradely transported via the vagal nerve from the gut to the brain to reach the SN ^7–9^. However, those studies did not show any evidence of neurodegeneration in the ventral mesencephalon. Recently, Dawson, Ko and colleagues have demonstrated that injection of preformed fibrils into the muscular layer of the pylorus and duodenum of mice leads to spread of pathological α-syn to the brain in a time-dependent manner, reaching the SN and causing dopaminergic cell loss and motor deficits ^10^.

To test the hypothesis that the gut-brain axis may be involved in the spreading of α-syn pathology under gut-related inflammation, we first analyzed the GI tract from DSS-treated rats (subchronic regime) and IBD human subjects to determine whether pathological α-syn is formed during the course of the disease. A previous study demonstrated increased expression of α-syn in colon samples from CD patients ^43^. Typical features associated with human UC are nitrosative and oxidative stress, known to trigger fibrillation of unmodified α-syn ^44, 45^. Our analysis demonstrated the presence of pathogenic P-α-syn in Auerbach’s and Meissner’s plexuses in both rat and human inflamed colon. Also, P-α-syn staining was observed in non-neuronal cells, probably associated with immune cells. Remarkably, these ENS locations matched with those found in many PD subjects, even at early stages ^19, 46, 47^. Overall, our data suggest the importance of gut inflammation in promoting pathogenic α-syn aggregation in the GI tract, a potential route towards the brain.

It is known that most distinguished features of PD are selective death of nigral dopaminergic neurons and the appearance of pathogenic α-syn aggregates. Consequently, we analyzed the integrity of the nigral dopaminergic system and α-syn aggregation in the ventral midbrain in a subchronic DSS rat model (28 days). Intriguingly, we found a loss of 40% of dopaminergic neurons in the right SN but no significance loss within the left SN. This asymmetry correlates with the innervation of the vagus nerve to the colon (right vagus nerve branch ^48, 49^) (Fig. 10). Interestingly, PD initially starts as a unilateral disorder, a common feature used for correct diagnosis of the disease, and not to be confounded by other conditions including essential tremor, multiple system atrophy and supranuclear palsy ^27, 50–52^. Subsequently, we analyzed the secondary structure of proteins including β-sheet rich α-syn amyloid fibrils typically associated with LB and Lewy neurites ^53^ by FTIR in the DSS subchronic model. This analysis confirmed the presence of insoluble α-syn aggregates and their preferential distribution in the right hemisphere of DSS-treated animals. These studies support the view of a bottom-up spread of the pathology. In order to further validate our data, a new cohort of vagotomized animals undergone the subchronic DSS treatment. We found pathological P-α-syn in the gut, but neither TH-cell degeneration nor α-syn aggregation in the ventral mesencephalon. These results support the important role of the vagus nerve in spreading pathogenic forms of α-syn from the inflamed gut to the brain. Interestingly, vagotomy also completely abrogated the long-term DSS-induced activation of key inflammatory markers in the ventral mesencephalon. Ventral midbrain neuroinflammation may be inherently associated to neurodegenerative events in response to DSS-induced gut inflammation since vagotomized animals showed both protection of the nigral dopaminergic system and absence of inflammation. In either case, these experiments support the involvement of the vagus nerve in spreading up PD-associated pathology. However, an important non-mutually exclusive feature of IBD and DSS-based models of UC is peripheral inflammation. It is long known that systemic inflammation causes microglial activation, cytokine production and selective death of nigral dopaminergic neurons in different experimental models^32, 54^. These events may have a significant contribution to dopaminergic pathology that needs to be further elucidated. Indeed, any condition accompanied by peripheral inflammation, including IBD, could present a risk factor for the significant neurodegeneration observed in PD ^55^. For instance, sustained peripheral inflammation may result in BBB dysfunction, which is critical to regulate extravasation of peripheral cytokines and immune cells, and thus, subsequent neuroinflammation.

**Figure 10.**
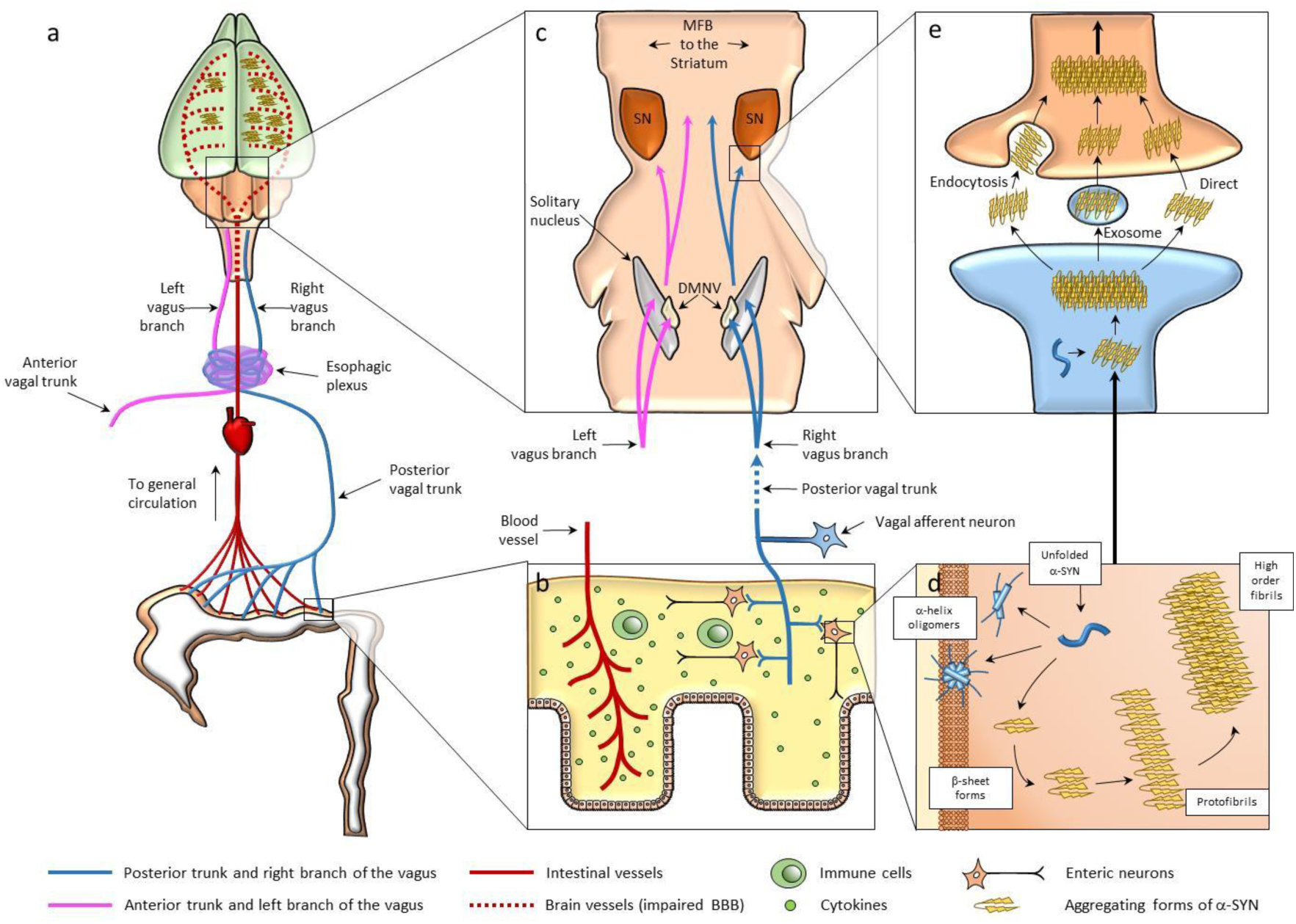
Schematic of proposed relationship between UC and the asymmetric death of dopaminergic neurons in the SN. Events in the gut can influence the brain in two ways: the humoral route and the neural route. The former consists of cytokines and other substances produced by enteric cells that are collected by the intestinal blood vessels and incorporated into the general circulation, eventually reaching the brain. The latter mainly comprises the multitude of nerve terminals and neurons distributed throughout the gut and eventually connecting with the vagus nerve. Although it is bilateral, the vagus nerve innervates the different organs of the body in an asymmetric way. The large intestine receives afferents from the posterior trunk of the vagus, which becomes the right branch once it passes through the diaphragm. The right and left branches mix incompletely in the esophageal plexus, so approximately 70% of the nerve connections from the large intestine project through the right branch of the nerve. Thus, in the rat DSS model of UC (**a**), the humoral pathway homogeneously distributes pro-inflammatory cytokines produced by immune cells in the large intestine (**b**), generating a pro-inflammatory environment in the brain (green shading) owing to alterations in the BBB (dashed red lines). At the same time, the right branch of the vagus nerve facilitates the arrival (mainly) to the right hemisphere of the brain (**c**) of elements that will contribute to asymmetric neurodegeneration, as explained below. In the large intestine, enteric neurons (**b**) use α-syn as part of the transmission system. Under inflammatory conditions, α-syn may adopt different conformations, including intrinsically disordered forms, forms rich in alpha helix and forms rich in beta sheet (**d**); the latter can form orderly oligomers of low order that accumulate in higher order structures such as protofibrils and fibrils. The neural pathway could transport these aggregates from one neuron to another (**e**), taking them from the intestine to the brain where they would form LB and Lewy neurites.

Consequently, we analyzed different features typically associated to peripheral inflammation including i) BBB integrity, ii) brain infiltration of immune cells, iii) microglia activation and iv) analysis of pro-inflammatory markers in brain and serum. We performed MRI studies in rats under DSS treatment and analyzed cerebrovascular functions and BBB integrity. Our results show that acute DSS induces a significant increase in rCBV together with a progressive increase in BBB permeability, in both the SN and striatum. Interestingly, in agreement with our results, it has been previously shown how an increase in rCBV may precede BBB permeability under neurodegenerative conditions^20, 56, 57^. In both cases, the observed changes likely reflect DSS-induced inflammatory effects on the vasculature. Furthermore, changes in BBB permeability were accompanied by a significant upregulation of ICAM-1 mRNA levels in the SN, a cell adhesion molecule that participates in the attachment of leukocytes to the vascular endothelium and its subsequent extravasation ^58^. Interestingly, this molecule has been suggested to sustain inflammation ^59^ and it is up-regulated in PD ^60, 61^. Indeed, our FACS, dot blot and confocal data suggested infiltration of peripheral monocytes and CD4^+^ and CD8^+^ lymphocytes at 2-4 days of DSS treatment, when the BBB was altered. Interestingly, infiltrating cytotoxic CD4^+^ and CD8^+^ lymphocytes, have been also observed in the inflamed SN of post-mortem human PD ^62–64^. Analysis of microglia revealed an early morphological activation in the ventral mesencephalon in parallel with the BBB alteration, also mimicking PD features ^65–67^.

Given the co-existence of gut and peripheral inflammation at the end of the subchronic DSS model and its relative short duration (28 days), we decided to extend the duration of the DSS regime (x3 cycles of DSS treatment) with a final washout phase of 14 days (chronic DSS model; 63 days). With this second model we wanted to study the long-term effects of α-syn pathology in gut and brain under the IBD-related systemic inflammation. We found that all analyzed inflammatory markers in the colon were not different from control animals. Analysis of α-syn pathology clearly revealed P-α-syn aggregates in the mucosal nerve fibers around the crypts, neuronal somas and nerve fibers of both the submucosa and the muscular external layer, a common pattern found in PD patients ^34, 35^.Further analysis of the ventral midbrain revealed the presence of α-syn inclusions within dopaminergic processes and, importantly, a significant loss of TH dopaminergic neurons in both hemispheres. Therefore, we may conclude that in our experimental models (subchronic and chronic DSS models), we clearly found lesions of the nigral dopaminergic system which started unilaterally (subchronic model) and then spread bilaterally (chronic model), mimicking the natural progression of PD and a long-lasting α-syn pathology.

In agreement with our findings, previous experimental models^10^ have supported the pathological α-syn spread from the gut to the brain, including the SN, and therefore, causing significant neurodegeneration. However, the clinical relevance of these experimental models is still to be fully determined. To shed light on this issue, we collected mesencephalic resections from eight different subjects affected by IBD: five cases of UC and three cases of CD, with ages ranging 37-65 years. We analyzed levels of α-syn phosphorylation at Ser129, which is considered to be a robust marker of α-syn inclusion pathology, and thioflavin-S staining where used to detect insoluble fibril formation ^68^. Hierarchical cluster analysis identified four well-defined neuronal conditions in IBD subjects: i) healthy melanized neurons lacking P-α-syn aggregation; ii) neurons with decreased amount of neuromelanin lacking P-α-syn aggregation; iii) neurons showing evidence of soluble P-α-syn inclusions with a clear displacement of neuromelanin to the periphery, suggesting that α-syn aggregates cause this displacement, and iv) highly depigmented neurons showing insoluble P-α-syn aggregates. To the best of our knowledge this is the first study demonstrating α-syn pathology within ventral midbrain dopaminergic neurons from patients earlier diagnosed with IBD and with an average age below 48 years. No signs of α-syn pathology were found in neurons from aged-matched controls, thus confirming our observations. These stages of neuronal changes resemble the different steps leading to LB formation ^69, 70^. In fact, typical features of the stages preceding LB appearance are poorly pigmented neurons containing α-syn cytoplasmic inclusions, similar to those reported here, including pale bodies ^69, 70^. Overall, analysis of the human samples strongly supports the association between IBD and pathogenic α-syn aggregates in the ventral mesencephalon, and reinforces the emerging association between gut disease and PD.

Our study adds to different pieces of evidence supporting the connection between IBD and PD. For instance, there is a genetic link between both diseases ^36^. Examples include variants of the caspase activating recruitment domain 15/nucleotide oligomerization domain 2 (CARD15/NOD2) gene ^71^, and LRRK2 ^72^. In addition, different large epidemiological studies have shown that IBD patients have an increased risk of suffering from PD ^14, 73–75^. Moreover, IBD patients who received anti-TNF therapy had a lower risk of developing PD when compared to IBD patients without therapy^74^. Given that systemic inflammation is a critical component of IBD pathogenesis ^76^, its contribution was argued as a plausible explanation for this protective effect ^74^. However, the demonstration of α-syn pathology in gut and brain of IBD patients supports the bottom-up pathology hypothesis and thus, the spreading of α-syn from submucosal neurons to the ventral midbrain, via the vagal preganglionic innervation of the gut, becomes plausible. The results of the current study are also in accord with other epidemiological studies suggesting that truncal vagotomy is associated with a decreased risk of PD ^60, 77, 78^. The fact that only a small subset of IBD patients develop PD suggest that gut inflammation is a miscellaneous of risk factors including i) bottom-up spreading of α-syn pathology, ii) systemic inflammation causing BBB disruption, iii) brain immune cell infiltration and iv) microglia activation. Overall, our study revitalizes the predicted importance of the GI tract in starting PD pathology and paves the way for different research strategies to define prodromal conditions and to identify novel early biomarkers of PD.

## METHODS

### Animals and treatments

Male and female albino Wistar rats (200–300 g) were used for these studies. Rats were kept at 22 ± 1°C and relative humidity (60%), a 12-h light–dark cycle and free access to food and water. A timeline of the treatment with DSS and points at which the different parameters have been measured is shown in Supplementary Fig. 13. The control group received tap water at the different time points analyzed. Two models of experimental UC, based on the oral administration of DSS (molecular weight 36-50 kDa; MP Biomedicals) at a concentration of 5% (w/v) in drinking water, were used. In the subchronic paradigm, both male and female rats received DSS for 1 week (with free access), following the method previously described by Okayasu ^79^. Subsequently, rats received tap water for 2 weeks and then continued for one more week with DSS (Supplementary Fig. 13a). Therefore, the full treatment duration was 28 days. In the chronic paradigm, male rats underwent 3 weeks of DSS separated from each other by 2 weeks on water, plus an additional washout period of 2 weeks on water after the third week with DSS (see Supplementary Fig. 13b). Therefore, the full treatment duration was 63 days. Effects of DSS administration, such as the existence of rectal bleeding, diarrhea, and weight loss, were evident at the end of the treatment. In addition to these clinical symptoms, histopathological alterations of the gut were similar to those seen in human UC. At the end of each treatment, animals were sacrificed. Other groups of male rats underwent the same concentration of DSS or water for two, three, four or seven days, depending on the technique used (Supplementary Fig. 13a). Both left and right hemispheres were analyzed separately. Experiments with animals were carried out in accordance with the Guidelines of the European Union Directive (2010/63/EU) and Spanish regulations (BOE 34/11370-421, 2013) for the use of laboratory animals; the study was approved by the Scientific Committee of the University of Seville.

### Bilateral sub-diaphragmatic vagotomy

Prior to surgery, rats were solid food-deprived overnight, whilst access to water was provided *ad libitum*. Rats were anesthetized by intraperitoneal injection of a mixture containing ketamine/medetomidine (0.5 and 40 mg/kg) and subjected to bilateral subdiaphragmatic vagotomy (n = 16). A midline abdominal incision (1 cm, caudal from the xiphisternum) was made along the linea alba. Tissue layers were retracted, and the liver and stomach carefully displaced to provide access to the esophagus. Both dorsal and ventral branches of the vagus nerve were exposed. Each vagal nerve trunk was isolated from the surrounding connective tissue and vasculature, ligated with surgical suture, and clearly cut at two levels. The in-between vagal segment was extracted to confirm total vagotomy. Finally, the abdominal muscles and the skin were sutured with surgical silk and wound clips, respectively. For sham-operated controls (n=16), the same surgical procedure was performed to expose the main vagal nerve trunks, but these were not ligated and excised. In each experimental condition, rats were treated with DSS (n =10) or tap water (n=6) 28 days after surgery, following the same protocol described above.

### Human tissue samples

Human samples of inflamed colon were obtained from subjects with UC (N=6). Healthy colon was obtained from subjects (control) who had undergone colon resection to remove a tumour (N=6) (Supplementary Table 1); these samples were taken from a healthy region away from the tumour. The age of the subjects ranged from 31 to 83 years old. Colon sections, embedded in paraffin, were obtained from University Hospital Virgen del Rocío (HUVR)-IBiS Biobank (Andalusian Public Health System Biobank and ISCIII-Red de Biobancos: ISCCII-PT13/0010/0056). This study was carried out in accordance with the principles of the Helsinki Declaration.Tissues were subjected to a pathological examination at the hospital to confirm the diagnosis of either UC or healthy colon.

Brain human tissues were obtained from subjects (age range: 37-59) who suffered (N=6) or not (N=6; control subjects) IBD and died from different causes (see Supplementary Table 2), kindly provided by brain banks from either HUVR-IBiS Biobank (reference number PT17/0015/0041) and the Oxford Brain Bank (OBB; reference number OBB 453). The region investigated for this study was the ventral mesencephalon. The brain sections were subjected to a neuropathological analysis according to routine procedures.

Human studies were approved by the corresponding ethic committees (authorization code: 2051-N-20). Informed consent was obtained from all subjects.

### Immunohistological evaluation

Animals used for immunohistochemistry of TH completed 28 or 63 days of treatment, whilst animals used for immunohistochemistry of Iba-1 completed 3 days and 28 days of DSS treatment, before being perfused through the heart under deep anesthesia (isoflurane) with saline150 to 200 ml and then 150– 200 ml of 4% paraformaldehyde in phosphate buffer, pH 7.4. After perfusion-fixation, brains were removed and then cryoprotected serially in sucrose in phosphate-buffered saline (PBS), pH 7.4, first in 10% sucrose for 24 h and then in 30% sucrose until sunk (2–5 days). Brains were then frozen in isopentane at -80°C. Incubations and washes were performed in Tris-buffered saline (TBS) or PBS, pH 7.4, and all work was performed at room temperature (RT). Primary and secondary antibodies used are listed in Supplementary Table 3. For immunohistochemistry of Iba-1 and TH, thaw-mounted coronal sections (20 µm) were cut on a cryostat at -20°C and mounted on gelatin-coated slides. Sections were washed and then treated with 0.3% hydrogen peroxide in methanol for 20 min and washed again.

For Iba-1 staining, sections were incubated in a solution containing TBS and 1% goat serum (Vector) for 60 min in a humid chamber. Slides were then drained and further incubated with the primary antibody in TBS containing 1% goat serum and 0.25% Triton-X-100 for 24 h. For TH staining, sections were incubated in a humid chamber with a solution containing PBS, 0.1% Triton-X-100, 1% bovine serum albumin (BSA; Sigma) and 2% goat serum for 1 h. Subsequently, slides were drained and further incubated with the primary antibody in a solution containing PBS, 0.5% Triton-X-100, 1% BSA and 2% goat serum for 12 h at 4°C in a humid chamber. For both TH and Iba-1 staining, sections were next incubated for 2 h with biotinylated IgG. The secondary antibody was diluted in TBS containing 0.25% Triton-X-100 (for Iba-1) or PBS containing 0.5% Triton X-100, 1% BSA and 2% goat serum (for TH), and its addition was preceded by three 10-min rinses. Sections were then incubated with VECTASTAIN^®^-Peroxidase solution (Vector; 1:100). The peroxidase was visualized with a standard diaminobenzidine/hydrogen reaction for 5 min. Some sections used for TH staining were counterstained with cresyl violet (Sigma). Slides were then mounted in DPX (BDH Laboratories).

For immunohistochemistry of α-syn in human samples, 5-μm thick tissue sections from paraffin blocks were dewaxed in xylene and rehydrated in a series of graded alcohols. Sections were treated with formic acid for 5-10 min, rinsed with water and then immunohistochemistry was performed automatically using the Benchmark ULTRA platform (Roche) with a primary antibody for α-syn (Novocastra N-ASYN, clone KM51, Roche; 1:25) and a secondary antibody. Slides were then mounted in DPX (BDH Laboratories,). Antigen retrieval was performed using EDTA buffer (pH 8.0).

For immunofluorescence, animals completed either 3, 28 or 63 days of treatment before being perfused. Immunofluorescence of Iba-1, CD4, CD8, α-syn, MAP-2 and TH in rat brains was performed on coronal brain sections (30 μm). Sections were permeabilized with 1% Triton X-100 in PBS for 1 h, blocked in 5% (w/v) BSA, 1% Triton X-100 in PBS for 1 h, and then incubated overnight at 4°C with the primary antibodies diluted in 1% (w/v) BSA, 1% Triton X-100 in PBS. The following day, sections were rinsed for 1 h in PBS containing 0.1% Triton X-100, incubated for 1 h with the corresponding secondary antibodies diluted in 1% (w/v) BSA, 0.1% Triton X-100 in PBS. Next, we wash the sections again with 0.1% Triton X-100 in PBS for 1h to remove the excess of secondary antibody. Sections were counterstained with Hoechst to visualize cell nuclei and were finally mounted in 50% glycerol.

For immunofluorescence of ubiquitin, α-syn, P-α-syn, UCHL-1 and thioflavin-S in rat and human colon and brain samples,5-μm thick tissue sections were cut from paraffin blocks, dewaxed in xylene and rehydrated in a series of graded alcohols. Brain sections were permeabilized with 1% Triton X-100 in PBS for 1 h, blocked in 5% (w/v) BSA, 1% Triton X-100 in PBS for 1 h, and then incubated overnight at 4°C with the primary antibodies diluted in 1% (w/v) BSA, 1% Triton X-100 in PBS. The following day, sections were rinsed for 1 h in PBS containing 0.1% Triton X-100, incubated for 1 h with the corresponding secondary antibodies diluted in 1% (w/v) BSA, 0.1% Triton X-100 in PBS, and then rinsed again with 0.1% Triton X-100 in PBS for 1 h. Colon sections were permeabilized by boiling with 0.01 M sodium citrate, pH 6, during 10 min. After blocking with 3% BSA, 3% fetal calf serum and 0.1% Triton X-100 in PBS, for 1 hour at RT, slides were incubated at 4°C overnight with the primary antibodies. Antibody binding was visualized with the appropriate fluorescent secondary antibodies. Misfolded proteins were detected by means of thioflavin-S (Sigma Aldrich; 1:1000) staining. Nuclei were stained with Hoechst 33258 (1:3000 dilution; Invitrogen), and sections were mounted in 50% glycerol.

### Immunohistochemistry data analysis

For the measurement of Iba-1 immunoreactivity, we used the AnalySIS imaging software (Soft Imaging System GmbH) coupled to a Polaroid DMC camera (Polaroid) attached to a light microscope (Leica Mikroskopie). Cells showing Iba-1 immunoreactivity were counted across five sections per animal, systematically distributed through the SN anterior–posterior axis. On each section, a systematic sampling of the area occupied by the Iba-1-positive cells was made from a random starting point, with a grid adjusted to count five fields per section. An unbiased counting frame of known area (40 µm x 25 µm = 1000 µm^2^) was superimposed on the tissue section image under a 40x objective. The different types of Iba-1-positive cells (displaying different shapes depending on their activation state) in the different treatment conditions (control, DSS 3 days and DSS 28 days) were counted as a whole. The entire z-dimension of each section (20-µm in all animals) was sampled; no guard zone was used; hence, the section thickness sampling fraction was 1.

The number of TH- and Nissl-positive neurons in the SN was estimated using a fractionator sampling design ^80^. Since unilateral tremor is one of the first motor symptoms of PD, left and right sides were analyzed separately. Counts were made at regular pre-determined intervals (x = 150 µm and y = 200 µm) within each section. The counted region had a thickness of 840 µm from plate number 38 to plate number 42 of the atlas of Paxinos and Watson ^81^. A systematic sampling of the area occupied by the SN was made from a random starting point. An unbiased counting frame of known area (40 µm x 25 µm = 1000 µm^2^) was superimposed on the tissue section image under a 60x objective. Therefore, the area sampling fraction was 1000 µm^2^/(150 µm x 200 µm) = 0.033. As above, the entire z-dimension of each section (20-µm in all animals) was sampled, and the section thickness sampling fraction was 1. In all animals, sections 100 µm apart were analyzed; thus, the fraction of sections sampled was 20/100 (0.20). The total number of TH- and Nissl-positive neurons in the SN was estimated by multiplying the number of neurons counted within the sample regions by the reciprocals of the area sampling fraction and the fraction of section sampled. To extrapolate the number of neurons of the entire structure from the volume sampled using the dissector, the software uses the Cavalieri method, which ensures that the final estimate is not biased.

For immunofluorescence quantification, images were acquired using an inverted ZEISS LSM 7 DUO confocal laser scanning microscope with exactly the same laser intensity and gain conditions. Images were quantified using Image-J software downloaded as a free software package from the public domain: http://rsb.info.nih.gov/ij/download.html), without any other treatment. All images are taken with the same Z, so they have the same distance. We also analyzed Hoechst staining to assess any variations in tissue permeability between brain sections. For P-α-syn, thioflavin-S and neuromelanin staining on brain sections, analyses were made only in TH-positive neurons.

### Dot blot

SN were placed in RIPA buffer (Millipore) diluted in PBS with a protease and phosphatase inhibitor cocktail (ThermoFisher) and homogenized by ultrasound on ice with ultrasonic processor UP100H (Hielscher, Germany). After centrifugation at low temperature and collection of the supernatant, protein content of the samples was estimated by the method of micro-Lowry using a standard of BSA (Fryer et al., 1986). 1µg of tissue homogenate was spotted in 1 µl volume aliquots onto 0.45 µm nitrocellulose membrane. On orbital shakers, membranes were blocked with blocking solution (5% skim milk powder (w/v) in 1% Tris-buffered saline with 1% Tween-20 (TBS-T1%) for 2 h at room temperature and then incubated with primary antibodies (Table 3) in blocking solution at 4°C overnight. Membranes were washed three times in TBS-T0.1% and incubated with horseradish peroxidase (HRP)-conjugated secondary antibodies in in blocking solution for 2 h at room temperature on an orbital shaker. Membranes were again rinsed three times in TBS-T0.1%. The Amersham™ Imager 600 was used to detect HRP-conjugated secondary antibodies using a chemiluminescent substrate (Biorad). Densitometry analysis was performed using ImageJ software.

### ELISA

Fresh peripheral blood was collected from animals of the different treatments in vacuum filled tubes and incubated at RT for 30 min to allow its coagulation. Then it was centrifuged at 2000 g for 15 min at 4 °C to obtain serum, which was stored at -40 °C until used. Serum TNF and IL-1β concentrations were determined by using a rat TNF alpha Uncoated ELISA and a rat IL-1β ELISA Kit (all from Invitrogen) following the manufacturer’s instructions. The plates were read on a Synergy HT multimodal plate reader (BioTek, Winooski, VT, USA) set to 450 nm. All conditions were assayed in duplicate.

### Measurement of DA, DOPAC and HVA

Analysis of striatal DA, and its metabolites DOPAC and HVA was performed by means of HPLC, equipped with a vwr-Hitachi Elite Lachrom L-2130 pump in conjunction with a glassy carbon electrode set at -550 mV (DECADE II, ANTEC), after 28 days of treatment with DSS. A Merck Lichrocart cartridge (125 mm x 4 mm) column filled with Lichrospher reverse-phase C_18_ 5 mm material was used. The mobile phase consisted of a mixture of 0.05 M of sodium acetate, 0.4 mM of 1-octanesulfonic acid, 0.3 mM of Na_2_EDTA and 70 ml methanol/l, adjusted to pH 4.1 with acetic acid. All reagents and water were HPLC grade. The flow rate was 1.0 ml/min. Measurement of all molecules in fresh tissue was performed according to the method previously described ^82^. Concentration of striatal DA, DOPAC and HVA was calculated with the aid of an eDAQPowerChrom 280 software.

### Magnetic Resonance Imaging (MRI)

All MRI was performed using a horizontal bore 9.4T magnet with a Varian DirectDrive™ (Agilent Technologies). Animals were anaesthetised with 1-2% isoflurane in 70% N_2_/30% O_2_ and a cannula positioned in the tail vein for gadolinium-DTPA injection. Animals were placed in a quadrature volume transmit-coil (i.d. 72 mm) with 4 phased-array surface receiver coil (i.d.: 40 mm, RAPID Biomedical GmbH), and positioned in the magnet. Body temperature and respiration were monitored throughout and maintained at ∼37°C and 50 breaths per min, respectively. Multi-parametric MRI was performed, including acquisition of T_1_-weighted images pre- and post-gadolinium-DTPA injection to assess BBB integrity, and T_2_-weighted images to determine macroscopic changes in tissue structure, as previously described ^20^. Briefly, T_2_-weighted images were acquired using a fast spin-echo sequence with a repetition time (TR) of 3.0 s and an effective echo time (TE) of 60 ms. Spin-echo T_1_-weighted images (TR = 500 ms; TE = 20 ms) were acquired both before and 5 min after intravenous injection of 100µl gadolinium-DTPA (Omniscan®, GE Healthcare). The matrix size and field of view were 256 x 256 and 3.5 cm x 3.5 cm, respectively. In all cases a multi-slice acquisition was used, spanning the two areas of interest (striatum and SN) with slice thickness of 1mm. Regions of interest (ROI) encompassing areas of visible signal change in both striatum and SN were drawn on both the T_2_-weighted and post-gadolinium-DTPA T_1_-weighted images, and the areas calculated.

To identify gadolinium enhancement in the targeted areas, ROI encompassing areas of both the SN and striatum were drawn on the image resulting from the subtraction of post-minus pre-contrast T_1_- weighted images.

### Regional cerebral blood volume (rCBV) map calculation and analysis

rCBV maps were generated from time-series images acquired during bolus injection of contrast agent and tracer kinetic analysis. A series of 40 FLASH gradient-echo images were acquired (TR 20 ms; TE 10 ms; flip angle 20°) at a rate of 1 image every 1.275 s, during which 100 µl of gadolinium-DTPA (Omniscan®, GE Healthcare) was injected via the tail vein cannula, over a 4 s period from image 8. The rCBV data were acquired with a 128 x 64 matrix and a 3.5 cm x 3.5 cm field of view to increase the temporal resolution. In all cases a multi-slice acquisition was used, spanning the two areas of interest (striatum and SN) with a slice thickness of 1 mm.

The rCBV maps were thresholded at a level that was equal to the mean signal intensity plus two standard deviations of the signal intensity of the prefrontal cortex (non-DSS-affected structure) as a reference. This approach allowed areas of increased rCBV to be more easily visualized and quantitative

An in-house ImageJ plugin was used to generate the maps, using the methods adapted from Smith et al. ^83^. Briefly, the relative concentration was calculated for the entire time series on a voxel-by-voxel basis using the following equation.

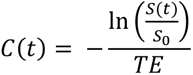

Where S(t) is the signal over time and S_0_ is the signal before the arrival of the bolus. The relative concentration was then integrated over time for each voxel after bolus arrival, to generate an rCBV map. As mentioned above, the rCBV of both the SN and striatum was normalized to the prefrontal cortex and these normalized values were compared. For each animal, we performed post – pre gadolinium-DTPA subtractions to generate a final map for each time point, and then compare absolute intensity throughout the time-course of the study.

### Quantification of peripheral immune cell infiltration by flow cytometry

Rats were anesthetized and perfused with saline solution until total clearance of peripheral blood, followed by decapitation following approved protocols. Brains were then removed and dissected; the SN was separated and placed on clean Petri dishes containing RPMI 1640 (Sigma). The tissue was then disaggregated by compression between two glass slides and clumps removed by passing the suspension through a 25G needle until disaggregation was complete. The suspension was then filtered through a 40 µm mesh filter and, subsequently, centrifuged for 10 min at 300 x g. The pellet was resuspended in 1 ml of PBS and aliquots of 100 µl were labeled with antibodies (all Becton Dickinson Bioscience) as shown in Supplementary Table 4. Finally, the cells were acquired in a FACSCanto II employing the FACSDiva software (Becton Dickinson Bioscience). Analysis was further conducted using the Flow Jo software program (FlowJo LLC; Supplementary Fig. 4).

### Real-time quantitative reverse transcription PCR

The left and right SN were dissected from each rat after either 3 or 28 days of treatment, snap frozen in liquid nitrogen and stored at −80°C. Total RNA was extracted using RNeasy® kit (Qiagen). cDNA was synthesized from 1 μg of total RNA using QuantiTect® reverse transcription kit (Qiagen) in 20 μl reaction volume, as described by the manufacturer. Real-time PCR was performed with iQ™SYBR® Green Supermix (Bio-Rad Laboratories), 0.4 μM primers and 1 μl cDNA. Controls were carried out without cDNA. Amplification was run in a Mastercycler® ep realplex (Eppendorf) thermal cycler at 94°C for 3 min followed by 35 cycles of 94°C for 30 s, 55°C for 45 s, and 72°C for 45 s, followed by a final elongation step at 72°C for 7 min. Following amplification, a melting curve analysis was performed by heating the reactions from 65 to 95°C in 1°C intervals while monitoring fluorescence. Analysis confirmed a single PCR product with the melting temperature. β-actin served as the reference gene and was used for sample normalization. The primer sequences for IL-1β, TNF, IL-6, ICAM-1, IL-10, arginase and β-actin are shown in Supplementary Table 5. The cycle at which each sample crossed a fluorescence threshold, Ct, was determined, and the triplicate values for each cDNA were averaged. Analyses of RT-qPCR were performed using a comparative Ct method integrated in a Bio-Rad System Software.

### Fourier transform infrared (FTIR) spectroscopy

FTIR spectra were recorded using Agilent Cary-FTIR microscope at SMIS beamline at SOLEIL synchrotron (France) and a Hyperion 3000 IR microscope (Bruker Scientific Instruments) coupled to a Tensor 27, which was used as the IR light source with 15x IR objective and focal plane array detector. The measuring range was 1000–4000 cm^−1^.Spectra collection was performed in transmission mode at 4 cm^−1^ resolution from 250 co-added scans, as described in Klementieva et al. ^84^. Background spectra were collected from a clean area of the same CaF_2_ window. All measurements were made at RT. Brain sections, 15 μm thick, were cut on a cryostat (Leica CM 1850 UV) and mounted onto CaF_2_ windows (Crystran).

Analysis of FTIR spectra was performed using OPUS software (Bruker). After atmospheric compensation, a linear baseline correction was applied from 1400 cm^−1^ to 2000 cm^−1^. Derivation of the spectra to the second order was used to increase the number of discriminative features and to eliminate the baseline contribution. Derivation of the spectra was achieved using a Savitsky−Golay algorithm with a 7- point filter and a polynomial order of 3. In FTIR spectroscopy, β-sheet structures can be distinguished based on the analysis of the amide I (1700–1600 cm^−1^). The level of β-aggregation of proteins in the tissue was studied in the left and right SN by calculating the peak intensity ratio between 1620 and 1630 cm^−1^, corresponding to β-sheet structures, and the maximum corresponding mainly to α-helical content at 1656 cm^−1^. An increase in the 1620–1630 cm^−1^component was considered a signature of β-sheet structures ^85^.

### Histological score of colon

For histological scoring, paraffin embedded sections of colon were stained with hematoxylin/eosin. The analysis was performed in a blinded fashion by a validated method ^86^. Colon damage was graded on a scale of 0–3 based on destruction of epithelium, dilatation of crypts, loss of goblet cells, inflammatory cell infiltrate, oedema and crypt abscesses.

### Statistics

Sample size for all experiments was calculated with the ENE 3.0 software (GlaxoSmithKline, Madrid, Spain). Groups were compared by one of the following tests when appropriate: the Kruskal-Wallis test for k independent measures, followed by the Dunn’s post hoc test; the Mann-Whitney U-test with a Bonferroni correction for post hoc pairwise comparisons; the Mann-Whitney test for 2 independent measures; the Friedman’s test for k related measures, with a Bonferroni correction for post hoc pairwise comparisons; or the Student-Newman-Keuls Method. The hierarchical cluster analysis, followed by a k-means method, was used for the analysis of human mesencephalic neurons (α=0.05) using the IBM SPSS 26 software. All measurements were taken from distinct samples, except peripheral cell infiltration data measured in a pool of tissue (Fig. 5d, e and f). The whiskers and boxes representations show quartile 1 (25%), quartile 2 (50%, median), quartile 3 (75%), as well as the 5^th^ and 95^th^ percentiles.

### Reporting summary

Further information on research design is available in the Nature Research Reporting Summary linked to this article.

## Data availability

The data that support the findings of this study are available from the corresponding author, upon reasonable request. Source data underlying Figs. 1, 2, 3, 4, 5, 7, 8 and 9 and Supplementary Figs. 2, 5, 6, 7, 8, 9 and 11 are available as a Source Data file.

## Acknowledgemensts

This work was supported by grants from the Spanish Ministerio de Economía y Competitividad (SAF2015-64171-R) and the Spanish Ministerio de Ciencia, Innovación y Universidades(RTI2018-098830-B-I00). AIRP and JLLG were funded by a grant of Spanish Ministerio de Salud (PI17/00828). MSS, SS and NRS were funded by Cancer Research UK (grant number C5255/A15935). VE was funded by a Marie Skłodowska-Curie Individual Fellowship. MJP, MDVC and PGM were supported by a grant from the Junta de Andalucía (CTS 5884) and thank Prof. Dr. A.A. Ilundáin for the support provided. The authors thank HUVR-IBiS Biobank (Andalusian Public Health System Biobank and ISCIII-Red de Biobancos) for providing the human samples and for the assessment and technical support provided. Images were obtained in the Centro de Investigación, Tecnología e Innovación de la Universidad de Sevilla (CITIUS).

## Author’s contributions

RMP and JLV designed the study and wrote the manuscript. RMP also carried out stereological analysis, PCR experiments, analyzed and interpreted the data. AMEO performed the animal treatments, perfusions and dissection of animal brains. She also carried out immunohistochemistry assays and was also involved in drafting and revising the manuscript. MSS, SS and VE performed MRI studies. MSS and RR performed and analyzed immunofluorescence assays from rat and human brains. ABS contributed to the immunohistochemistry assays, PCR and sample preparation for FTIR. AIRP performed bilateral sub-diaphragmatic vagotomies. MARC helped with the immunofluorescence and immunohistochemistry techniques. JGR helped with the PCR experiments. MS performed the HPLC assay. AEC, MDVC, PGM and MJP performed immunohistological assays from human and rat colons. MJP also contributed to the analysis and interpretation of results. OK carried out the FTIR study. MJOM and MSS performed the flow-cytometry assay. ER did the neuropathology analysis of the human brains. AM contributed to the design and conceiving of the manuscript. AJH contributed to statistical study, analysis and representation of data and writing of the manuscript. JLLG, TD and NRS were involved in critical revision of the manuscript. All authors discussed the results and commented on or edited the manuscript. All authors read and approved the final manuscript.

---

**Supplementary figure 1.**
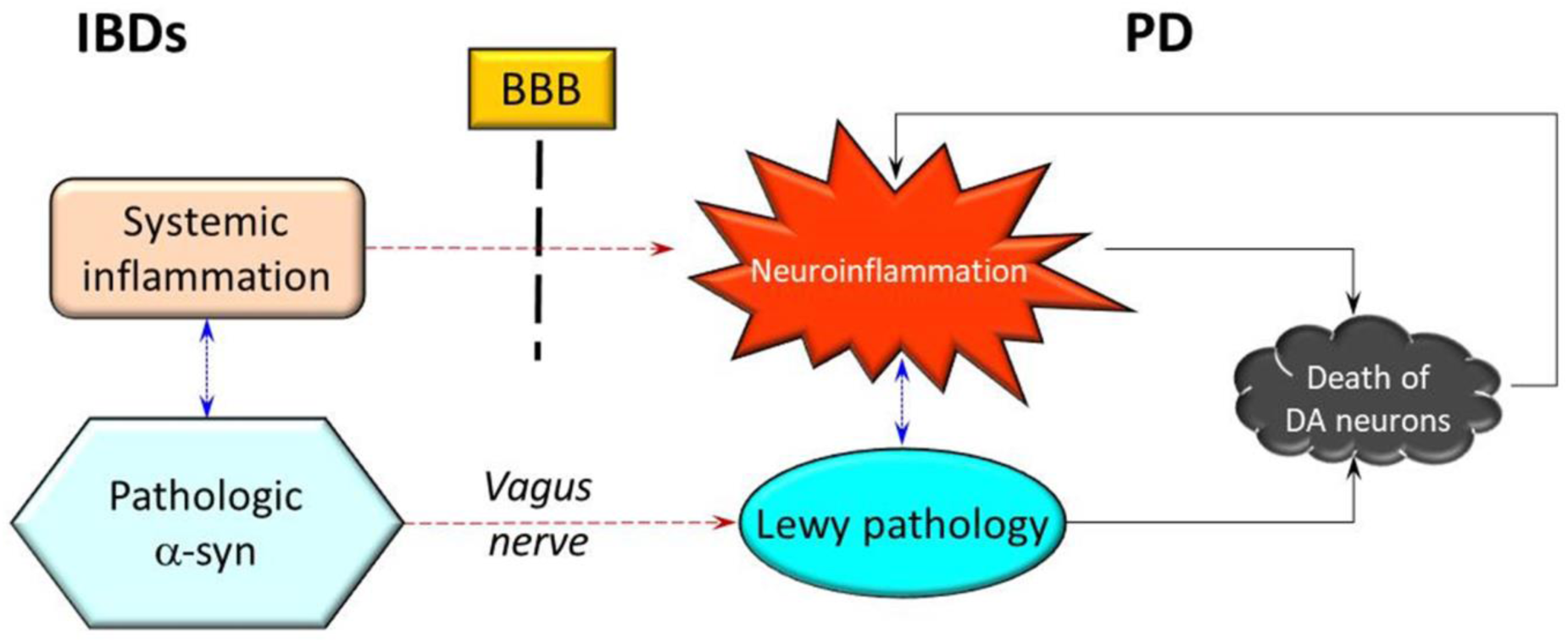
Connections between peripheral and brain inflammation and the role of α-syn. IBDs in the periphery and PD in the CNS may have parallel lives. Inflammation and alteration of -synuclein (-syn) might reinforce each other (blue dotted arrows) and may cause neuronal death; in turn, peripheral phenomena can influence the central ones (red dashed arrows) through a weakened BBB (inflammation) and through the vagus nerve (spread of -syn).

**Supplementary figure 2.**
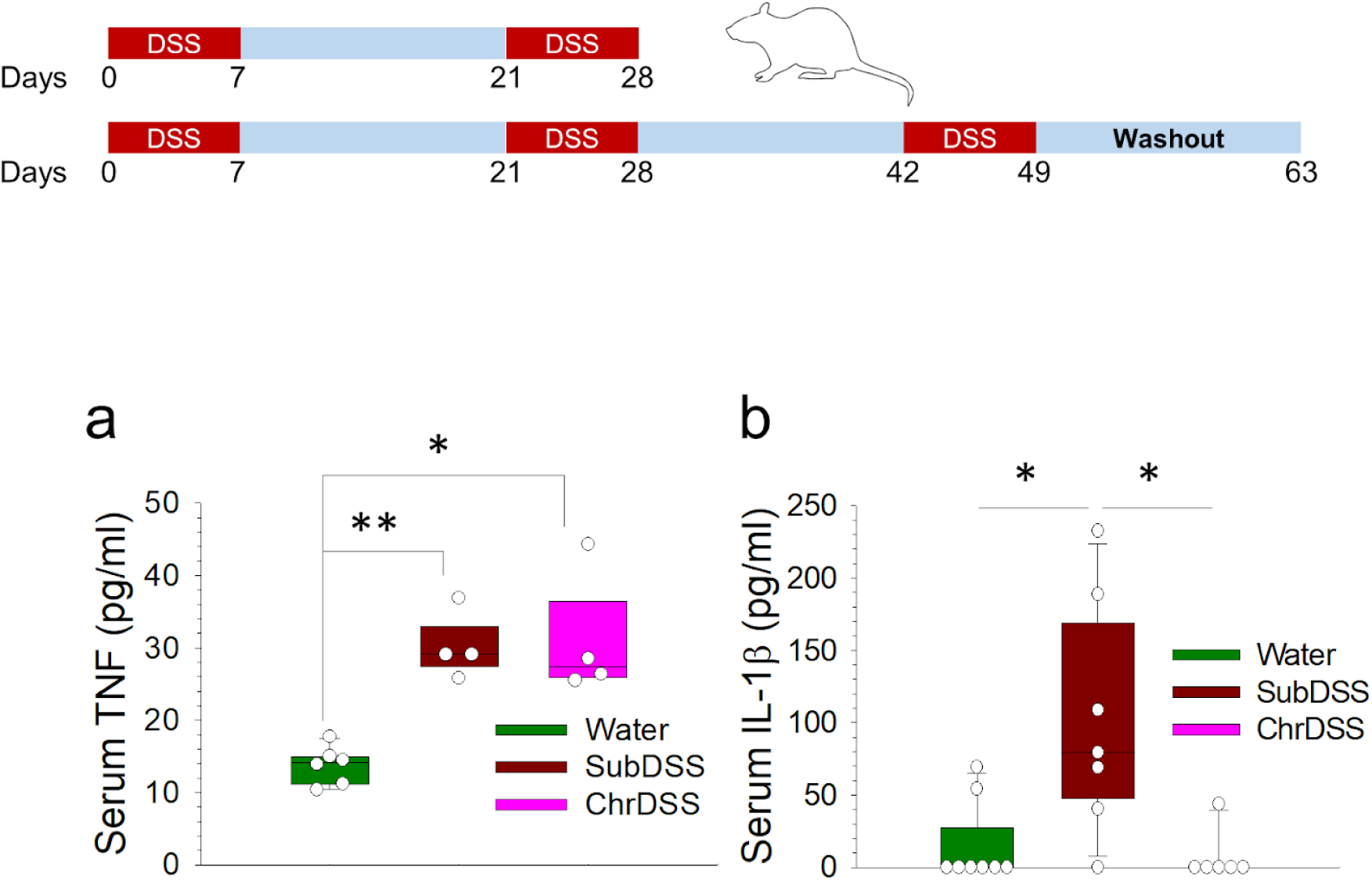
Effect of DSS treatments in peripheral inflammation. Serum levels of TNF (**a**) and IL-1β (**b**) were measure by ELISA. Results are expressed as pg/ml of the analyzed protein. N=4-8. Statistical analysis: Kruskal-Wallis test for k independent measures followed by the Dunn’s post hoc test for pairwise comparisons with the Bonferroni’s correction, with α=0.05: *, p<0.05, **, p< 0.01. Source data are provided as a Source Data file.

**Supplementary figure 3.**
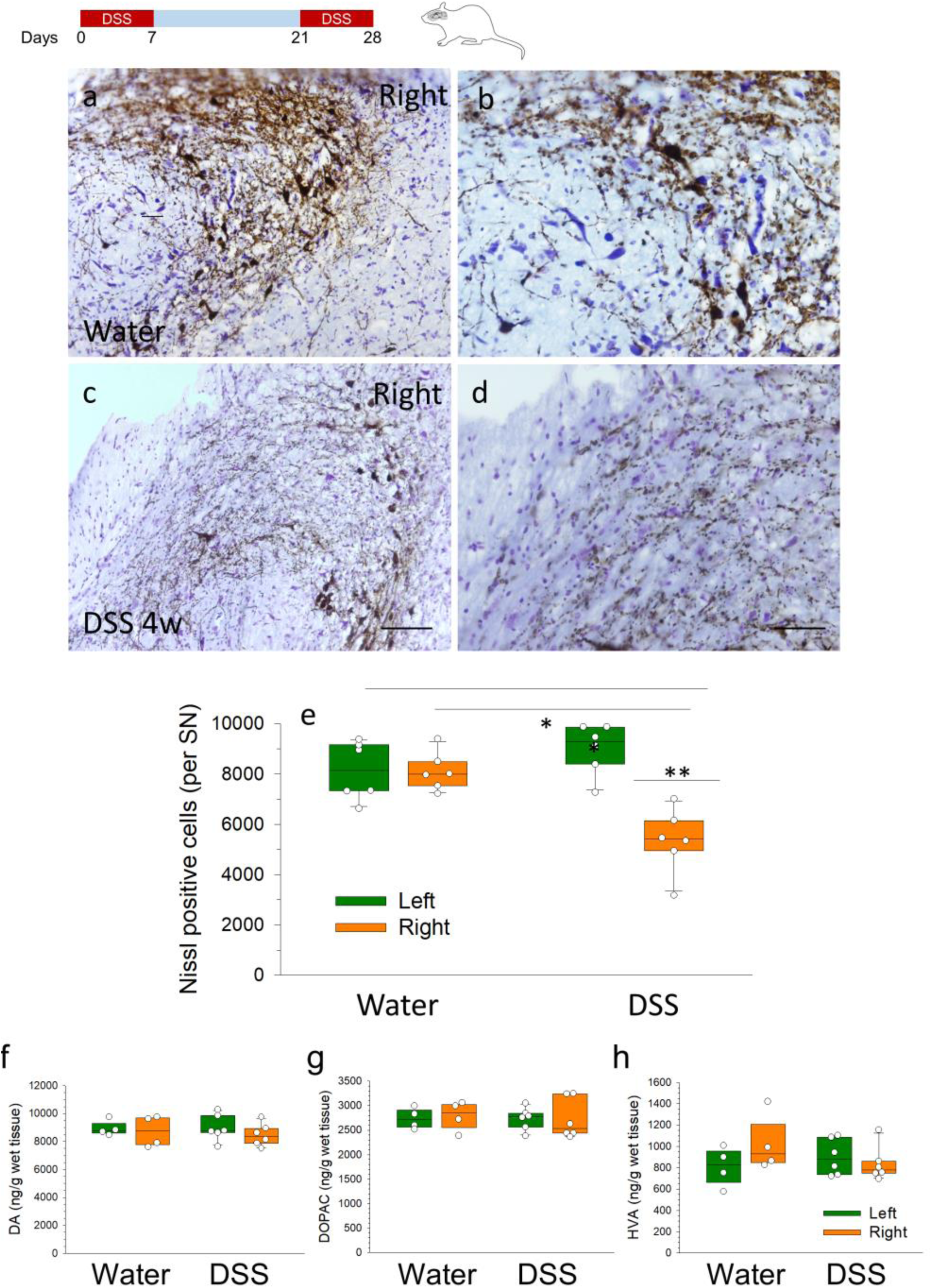
Nissl staining of the SN and DA, DOPAC and HVA measurements in striatum of control and DSS-treated rats. Representative photographs of the right SN of control (**a**) and DSS-treated animals (**c**).**b** and **d** show a magnification of panels a and c, respectively. A decrease in the number of Nissl-positive cells in the right side of the DSS-treated animals is observed. Scale bars: 125 µm (a and c) and 50 µm (b and d). (**e**) Quantification of Nissl-positive cells in the SN of rats. Results are expressed as number of Nissl-positive neurons. N=6 for all the groups. Statistical analysis: Kruskal-Wallis test for k independent measures followed by the Dunn’s post hoc test for pairwise comparisons with the Bonferroni’s correction, with α=0.05: *, p<0.05, **, p< 0.01. Source data are provided as a Source Data file. Amounts of dopamine (**f**) and its metabolites DOPAC (**g**) and HVA (**h**) in the striatum of control (water) and DSS-treated rats after 28 days of treatment. Results are nanograms per gram of wet tissue. N=4 for control and N=6 for DSS groups. Statistical analysis: Friedman’s test for k related measures followed by post hoc test for pairwise comparisons with the Bonferroni’s correction. No significant changes were found. Source data are provided as a Source Data file.

**Supplementary figure 4.**
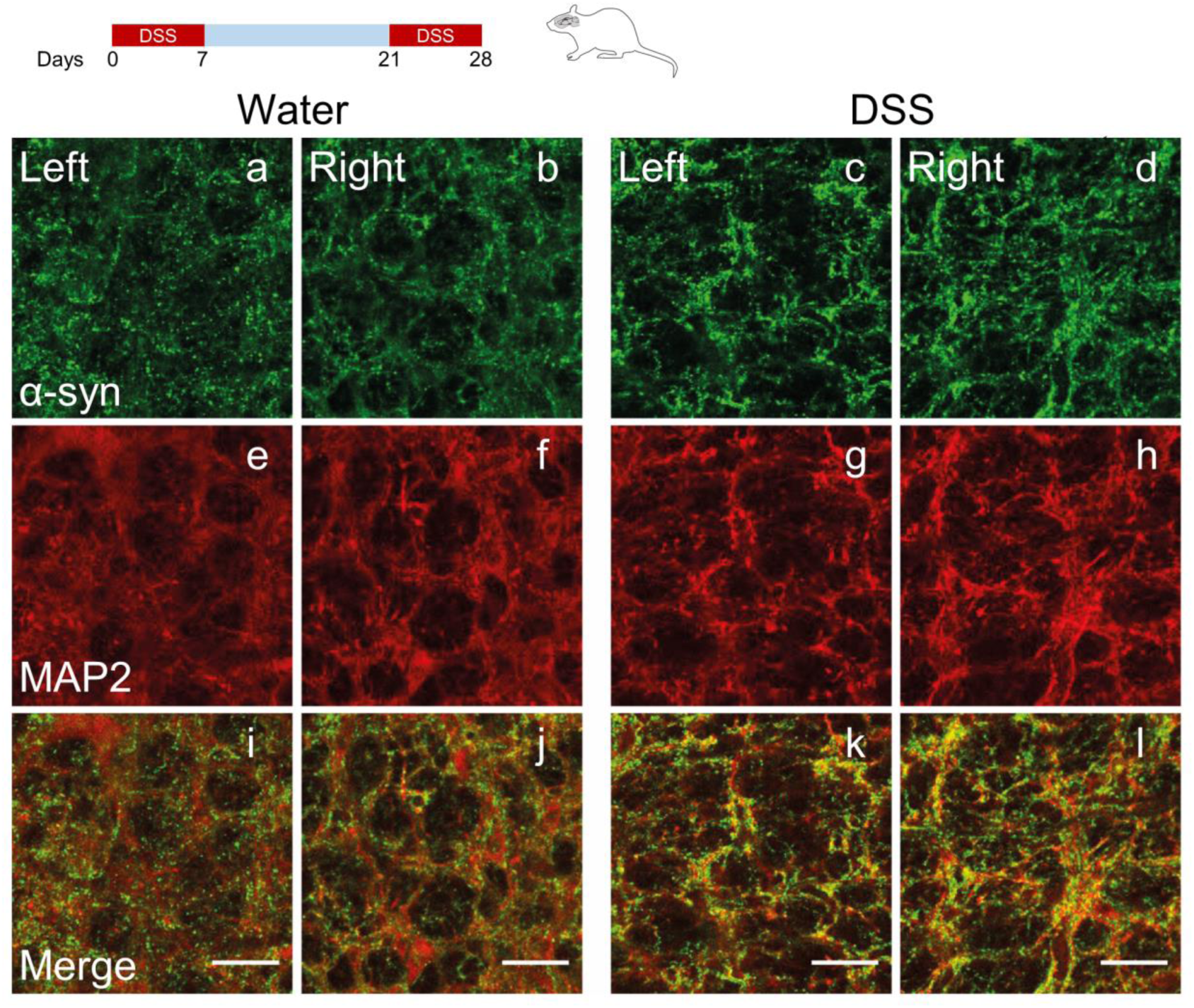
Immunofluorescence of α-syn and MAP-2 in the DMNV of control and DSS-treated animals. Representative microphotographs of immunofluorescence for α-syn (green) in control (**a**, left side and **b** right side) and DSS-treated rats (**c** left side and **d** right side), and for MAP-2 in control (**e**, left side and **f** right side) and DSS-treated rats (**g** left side and **h** right side). **i-l** correspond to merge of all images showed. Note the increment of α-syn intensity in the brain slides from DSS-treated rats in comparison with controls. Scale bars: 50 µm.

**Supplementary figure 5.**
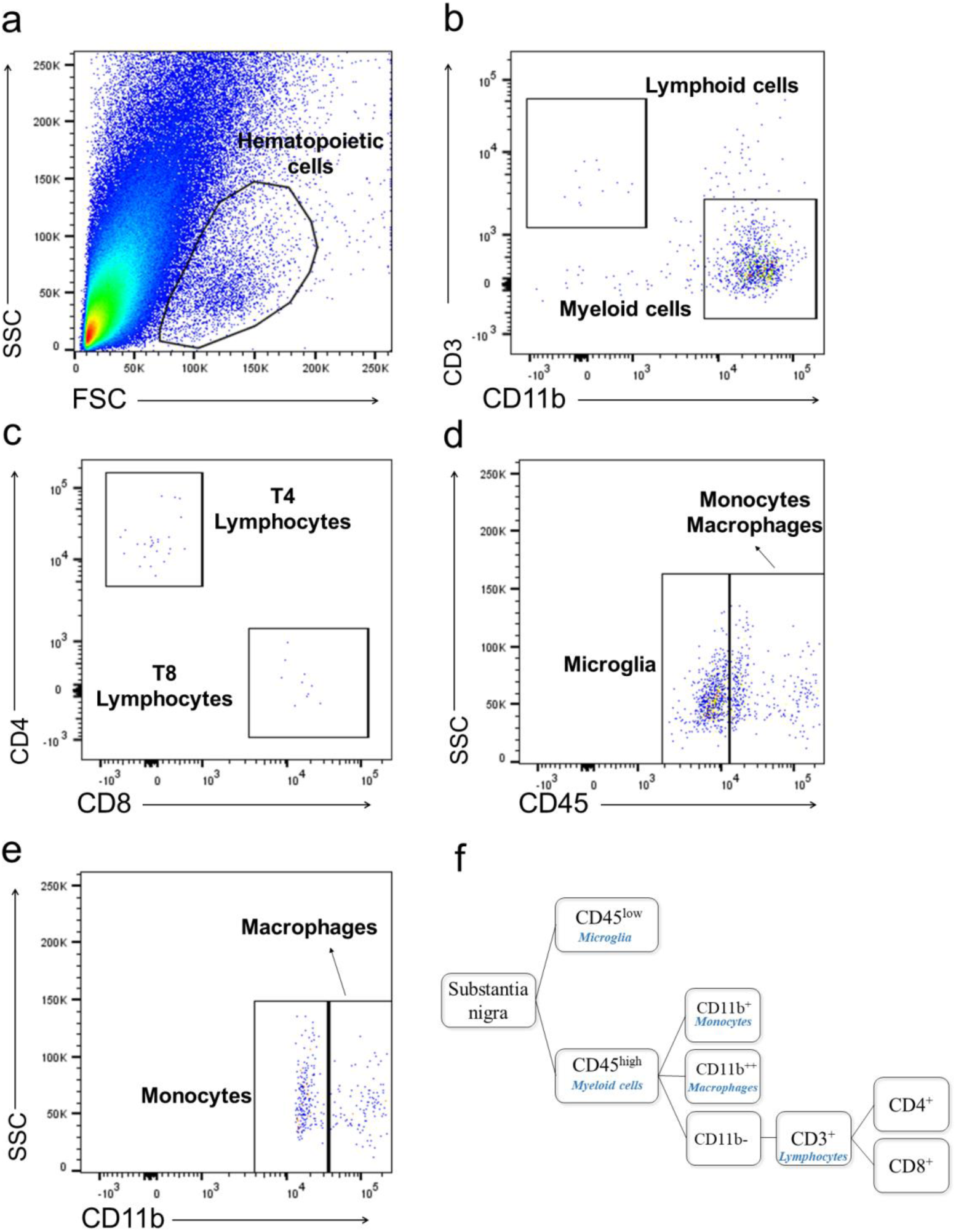
Infiltration of peripheral immune cells in an LPS injection model of BBB disruption. FACS analysis of SN tissue 48 h after LPS injection. (**a**) Forward and side scatter plot showing localization of hematopoietic cells. (**b**) After selecting CD45-positive cells, lymphoid and myeloid cells were subgated based on the expression of CD3 and CD11b, respectively. (**c**) T lymphocytes (CD11b^-^) were divided into T4 and T8 lymphocytes based on the expression of CD4 and CD8, respectively. (**d**, **e**) Monocytes, macrophages and microglia were characterized based on the expression of CD45 (d) and CD11b (e). Monocytes: CD45high CD11b^+^. Macrophages: CD45high CD11b^++^. Microglia: CD45low, CD11b^+^. (**f**) Representation of the gating strategy.

**Supplementary figure 6.**
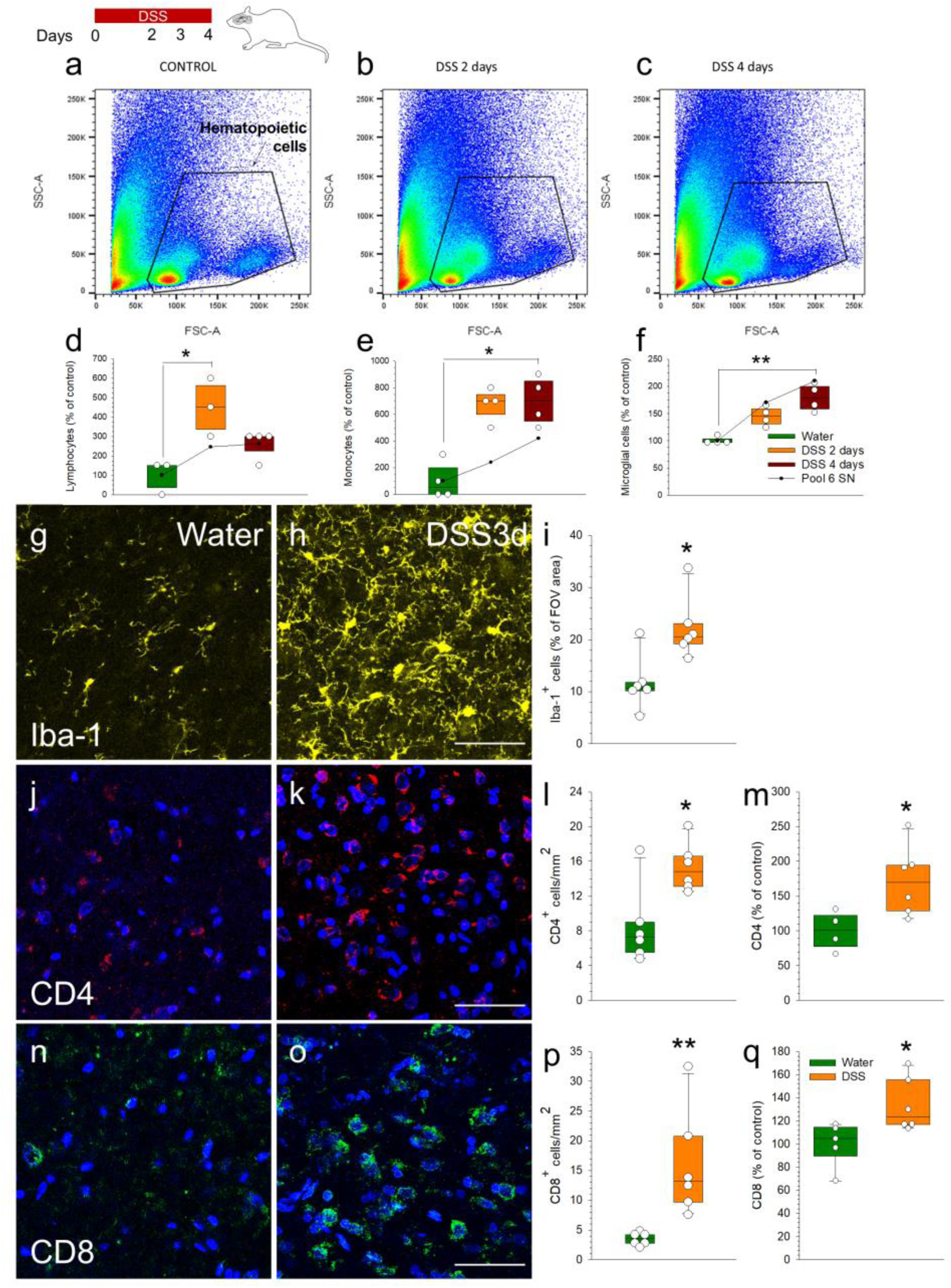
DSS causes infiltration of peripheral cells into the SN. FACS analysis in the SN of control animals (**a**) and animals after 2 (**b**) or 4 days (**c**) of DSS treatment. Forward and side scatters show the abundance of hematopoietic cells. Graphs show the percentage of lymphocytes (**d**), monocytes (**e**) and microglial cells (**f**) in the tissue sample. The results shown in the boxes and whiskers representations are expressed as percentage of control from individual SN (N=3 for the control and DSS 2 days groups of lymphocytes; N=4 for the rest of groups). The line and scatter plots represent a pool of N=6 SN. Peripheral cell infiltration was confirmed by immunofluorescence of Iba-1 for microglial cells (**g**, **h** and**i**), and of CD4^+^ (**j**, **k, l** and **m**) and CD8^+^ (**n,o, p and q**) for lymphocytes. Scale bar: 50 µm Results are expressed as percentage of microglial area per field of view (FOV), and number of CD4^+^ and CD8^+^ lymphocytes per mm^2^, respectively. Panels **m** and **q** show dot blot analysis as percentage of control. Statistical analysis for panels d-f: Kruskal-Wallis test for k independent measures followed by the Dunn’s post hoc test for pairwise comparisons with a Bonferroni correction, with α=0.05. Statistical analysis for panels I, l, m, p and q; Mann-Whitney U test for independent samples, with α=0.05; *, p< 0.05 and **, p< 0.01 comparing the DSS with the control group. Source data are provided as a Source Data file.

**Supplementary figure 7.**
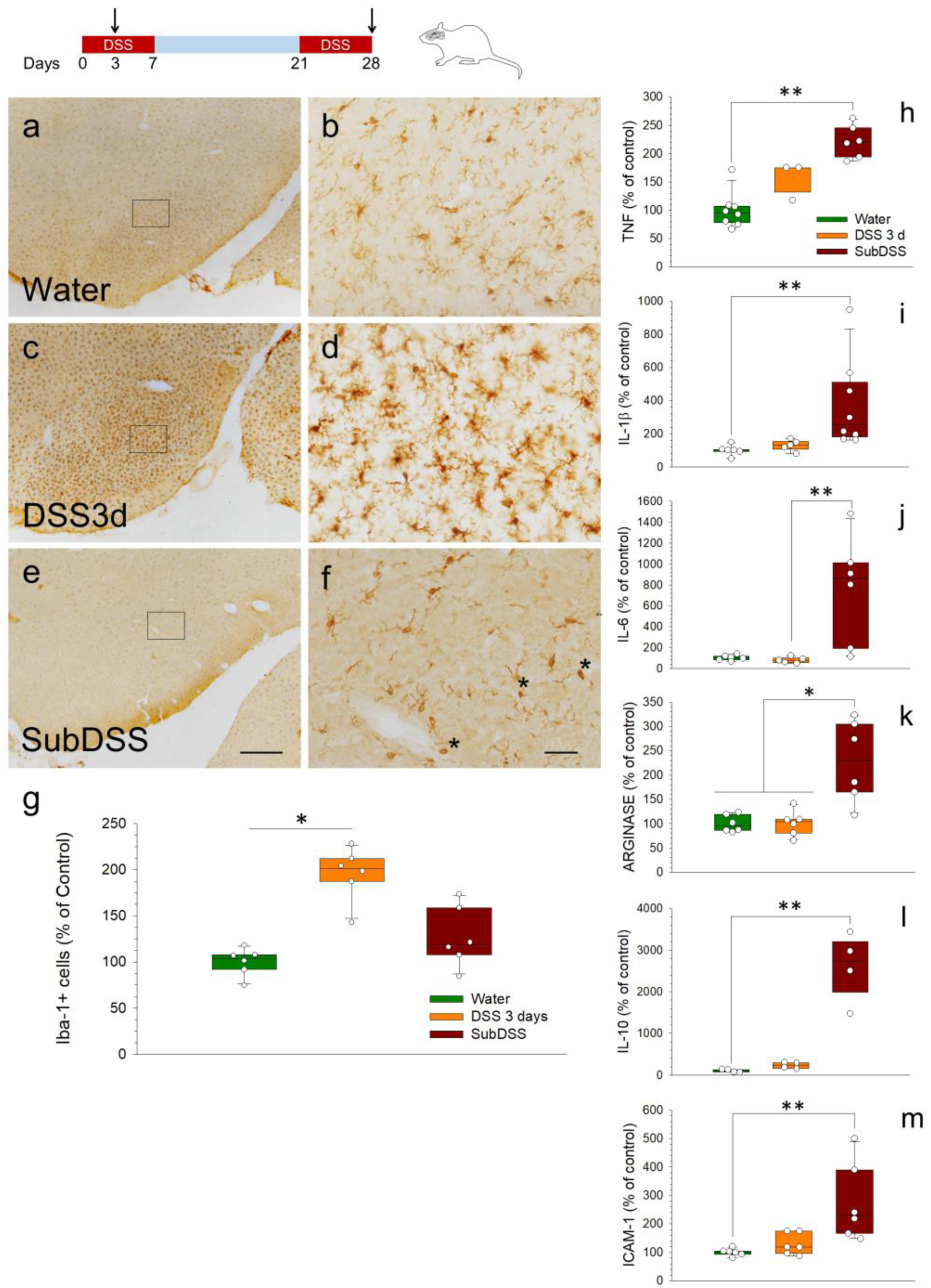
Effect of the DSS treatment on the activation of microglia in the ventral mesencephalon of male rats at both cellular and molecular levels. Coronal sections showing Iba-1 immunoreactivity in the SN. (**a, b**) Control animals. (**c, d**) Animals after 3 days of DSS. Note that DSS treatment increased microglial activation after 3 days. (**e, f**) Animals after 28 days of DSS. Asterisks point to round phagocytic cells. Scale bars: a, c and e, 500 μm; b, d and f, 50 μm. Panels b, d and f show a magnification of the boxes in the corresponding left panels. (**g**) Quantification of Iba-1-positive cells in the SN of rats. Results are expressed as percentage of control; N=6 for all groups. Effect of DSS on the expression of TNF (**h**), IL-1β (**i**), IL-6 (**j**), arginase (**k**), IL-10 (**l**) and ICAM-1 (**m**) mRNAs. mRNA expression in SN after 3 or 28days of DSS intake was quantified by RT-qPCR. No significant changes were found after 3 days of treatment with DSS. Expression levels of TNF, IL-1β, IL-6, arginase, IL-10 and ICAM-1 increased after 28 days of DSS. Results are expressed as percentage of control values; N=3-8. Abbreviations: DSS3d, 3 days of DSS; ChrDSS, 28 days of DSS. Statistical analysis: Kruskal-Wallis test for k independent measures followed by the Dunn’s post hoc test for pairwise comparisons with the Bonferroni’s correction, with α=0.05. *, p< 0.05 and **, p< 0.01, compared with the control group. Source data are provided as a Source Data file.

**Supplementary figure 8.**
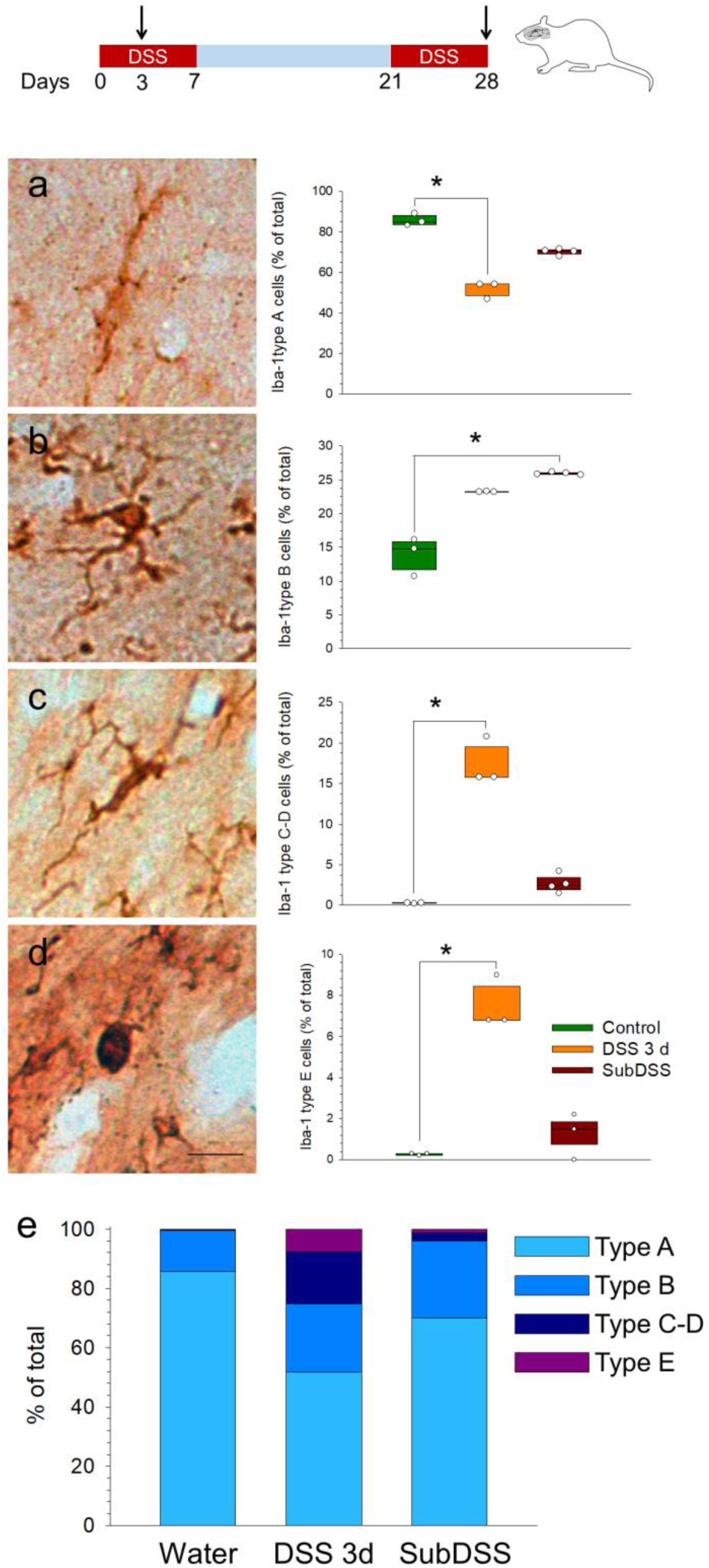
Change of microglial morphology along the activation process induced by the DSS treatments in rats. Microglial cells were counted separately according to their morphology. (**a**) Quiescent microglia (type A) has small cell bodies, fine cytoplasmic ramifications, and low to moderate Iba-1 expression. (**b**) Early-stage activation (type B) is characterized by increased ramification of cytoplasmic processes and cell size, and enhanced Iba-1 labeling. This is followed by further thickening of processes and retraction of the finest ones (**c**), as well as increased cell body size and Iba-1 expression (type C-D). (**d**) Near the completion of the microglia activation process, amoeboid cells may be observed (type E), showing complete retraction of cytoplasmic processes with maintained high levels of Iba-1 expression. Results are expressed as percentage of total cells; N=3 for control and DSS 3 days groups; N=4 for DSS 28 days group. Statistical analysis: Kruskal-Wallis test for k independent measures followed by the Dunn’s post hoc test for pairwise comparisons with the Bonferroni’s correction, with α=0.05. *, p<0.05. Scale bar: 20 µm. (**e**) Percentage of the different microglial morphologies in the control and DSS groups. Source data are provided as a Source Data file.

**Supplementary figure 9.**
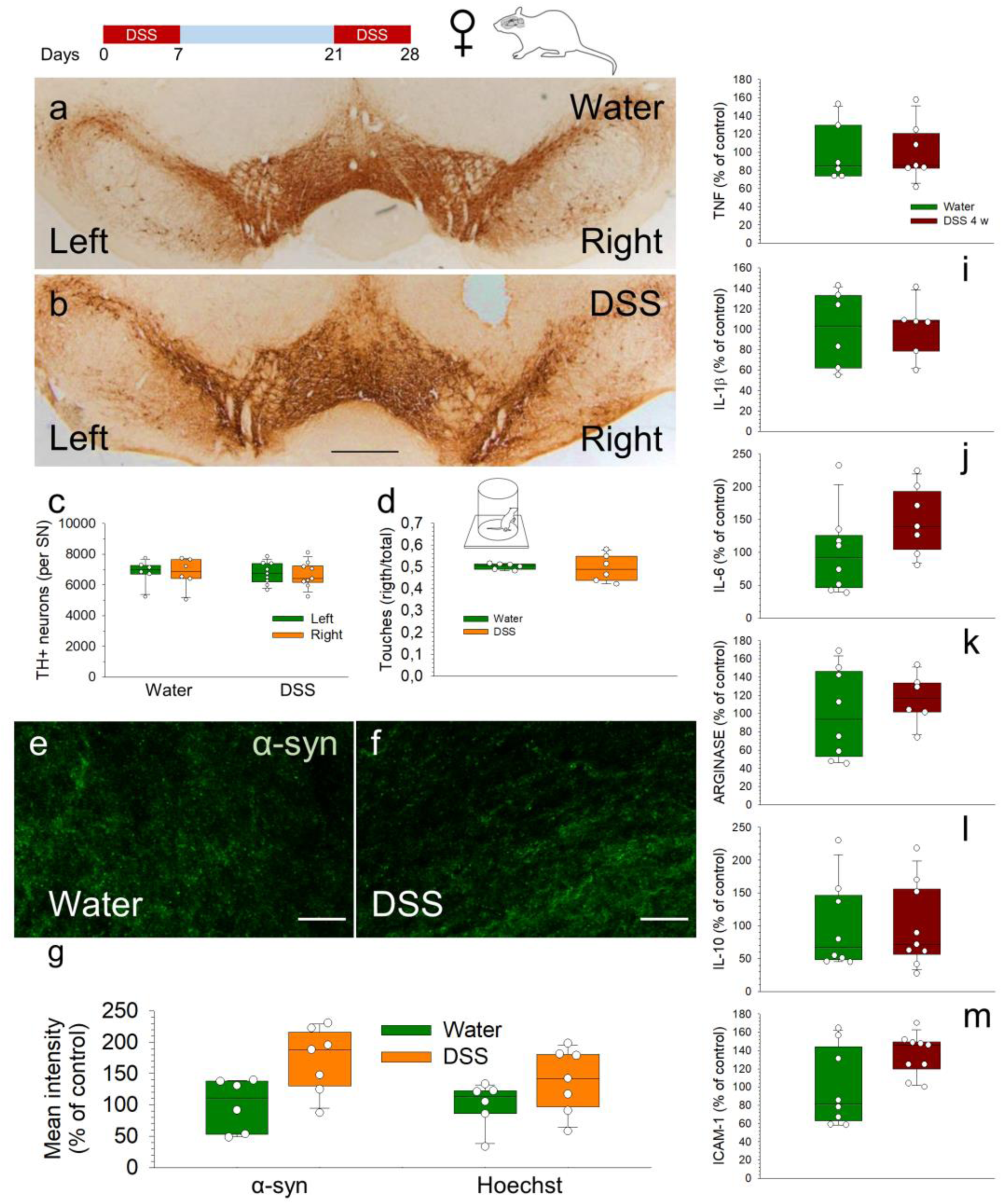
Effects induced by the subchronic treatment with DSS in female rats. Coronal sections of TH immunohistochemistry showing the whole SN from control (water, **a**) and DSS treated (**b**) female rats. Scale bar: 1 mm (a and b). (**c**) Quantification of TH-positive cells in the SN of rats. Results are expressed as number of TH-positive neurons per SN. N=6-9; no significant difference was found. (**d**) Quantification of number of touches with the right and left paw (Cylinder test). N=6 for each group; results are expressed as ratio of touches (right/total); no significant differences were found. Immunofluorescence shows no difference of -syn in the SN of control (**e**) and DSS-treated female animals (**f**). Scale bar: 30 µm (e and f). (**g**) Quantification of the mean fluorescence intensity of -syn and Hoechst intensity in the SN of female rats. Results are expressed as percentage of control values for each parameter. N=6-7. No significant differences were found. Effect of DSS on the expression of TNF (**h**), IL-1β (**i**), IL-6 (**j**), arginase (**k**), IL-10 (**l**) and ICAM-1 (**m**) mRNAs in female rats. mRNAs expression in SN after 4 weeks of DSS intake was quantified by RT-qPCR. No significant changes were found. Results are expressed as percentage of control values; N=6-9. Statistical analysis: Kruskal-Wallis test for k independent measures followed by the Dunn’s post hoc test for pairwise comparisons with the Bonferroni’s correction was used for statistical analysis in all the parameters measured except the cylinder test that was analyzed by the Mann-Whitney test, with α=0.05. Source data are provided as a Source Data file.

**Supplementary figure 10.**
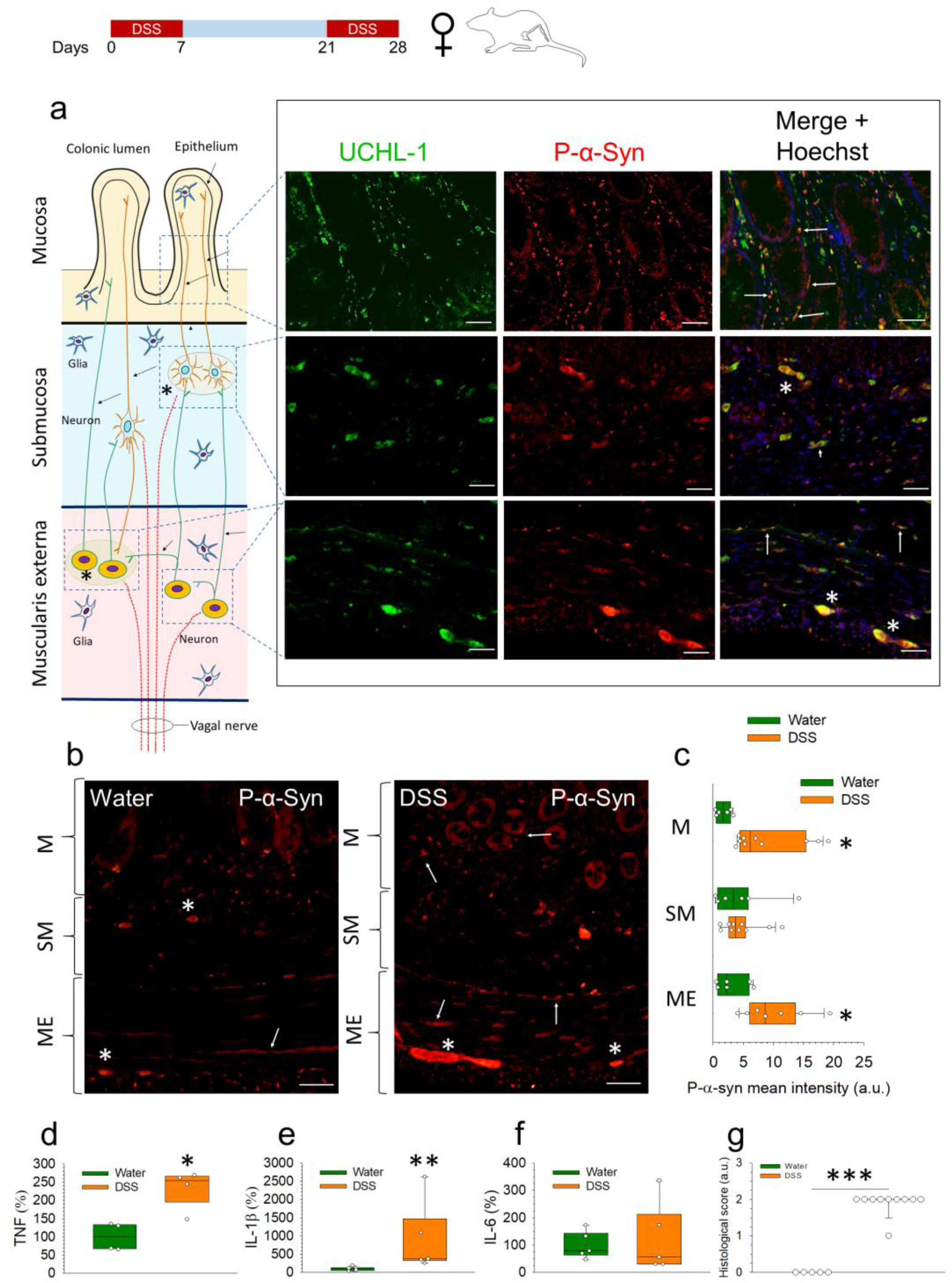
Immunolocalization of P-α-syn and measurement of pro-inflammatory cytokines in the colon of female rats under subchronic treatment with DSS. (**a**) Schematic cross-section of the colon wall illustrating the interconnected enteric plexuses, which contain neurons projecting to the mucosa, and vagal fibers. Co-localization of UCHL-1 and P-α-syn in the nerve fibers (arrows) and ganglia (asterisks) of the colon layers of a DSS-treated female rat. (**b**) Representative photographs of P-α-syn staining in the mucosa (M), submucosa (SM) and muscularis externa (ME) from colon of control (water) and DSS-treated female rats. Neuronal structures are indicated as in (a). Scale bars: 20 µm (a) and 50 µm (b). (**c**) Quantification of P-α-syn staining fluorescence intensity (as arbitrary units) measured in the mucosa (M), submucosa (SM) and muscularis externa (ME); N=6-7. (**d**, **e**, **f**) mRNA expression of TNF, IL-1β and IL-6 quantified by RT-qPCR in the colon of control (water) and DSS-treated female rats; N=4-5. (**g**) Histological score of the colon of control (water) and DSS-treated female rats; N=5-7. Statistical analysis: Mann-Whitney U test for independent samples, with α=0.05; *, p< 0.05, **, p< 0.01 and ***, p < 0.001, comparing the DSS-treated with the control (water) group. Source data are provided as a Source Data file.

**Supplementary Figure 11.**
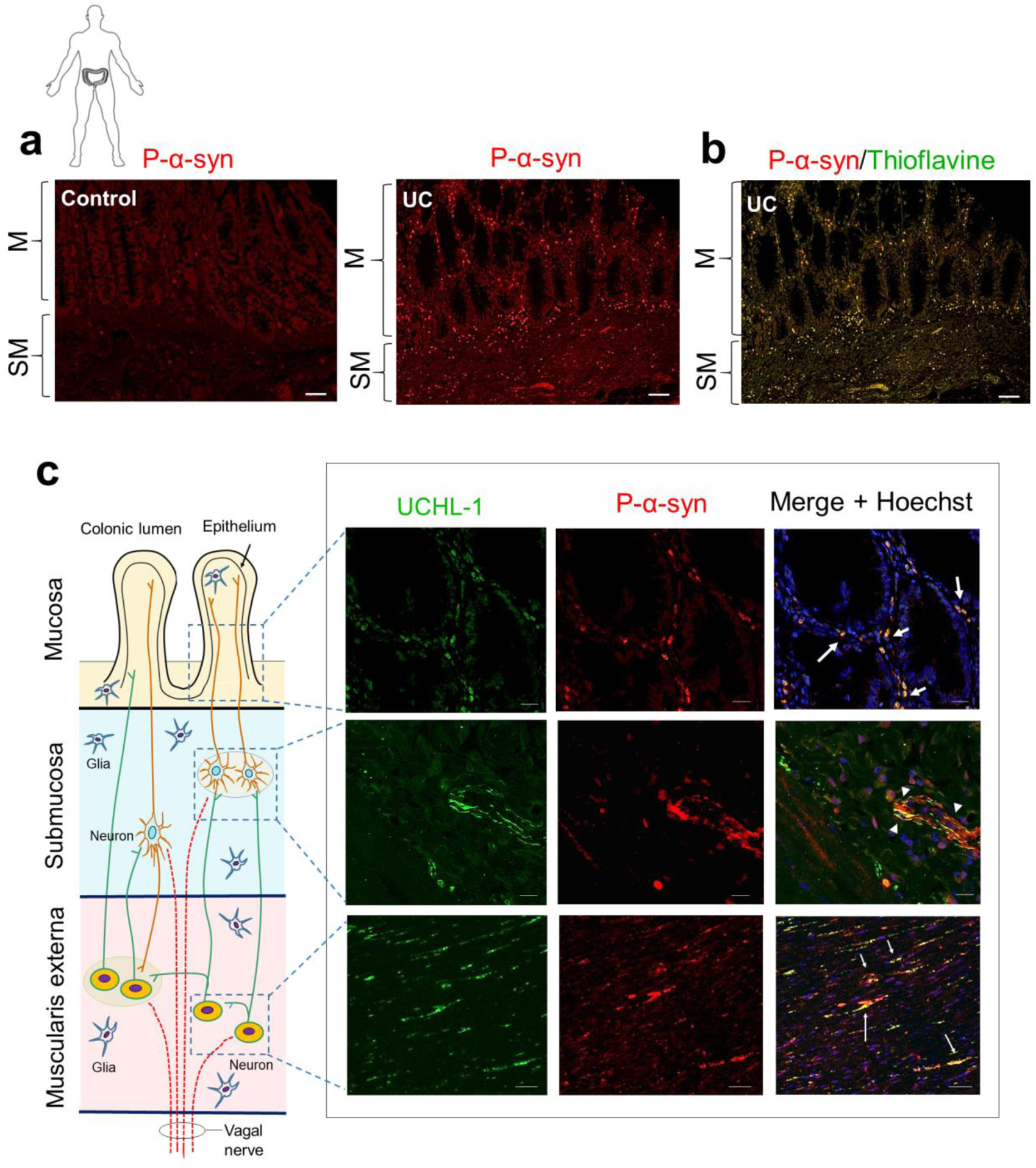
Immunolocalization of P-α-syn in the colon of elderly subjects with UC. (**a**) Representative photographs of P-α-syn staining in the mucosa (M) and submucosa (SM) from colon of elderly subjects (controls and with UC). (**b**) Co-localization of P-α-syn and thioflavin-S in the mucosal nerve fibers and submucosal plexus of an elderly patient with UC. (**c**) Schematic cross-section of the colon wall illustrating the interconnected enteric plexuses, which contain neurons projecting to the mucosa, and vagal fibers. Co-localization of P-α-syn and UCHL-1 in the mucosal nerve fibers around a crypt (large arrows), in the submucosal nerve fibers and ganglia (arrowheads), and in the neuronal somas and nerve fibers (small arrows) of the muscular layer of an elderly patient with UC. Scale bars: 50 μm (a, b) and 20 μm (c). Images are representative of N=6 assays.

**Supplementary Figure 12.**
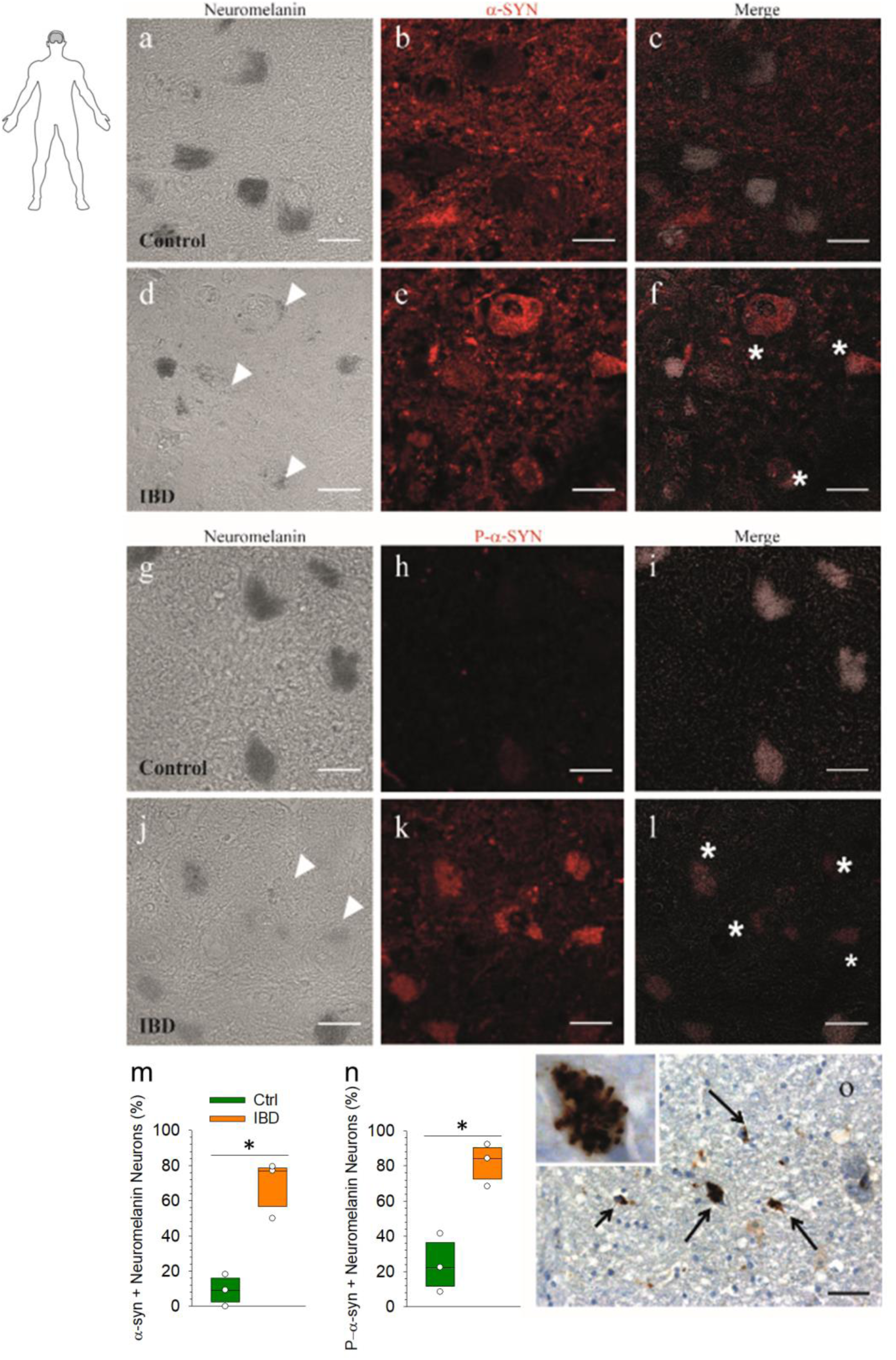
Accumulation of α-syn and P-α-syn in SN brain slides from IBD subjects. Representative confocal micrograph images of SN brain slides from a control case and an IBD patient. Left panels show neuromelanin-positive neurons (bright field; **a**, **d**, **g** and **j**); central panels show immunolabelling of α-syn (red; **b** and **e**) and P-α-syn (red; **h** and **k**); right panels show merge of both images (**c**, **f**, **i** and **l**); asterisks indicate colocalization; arrowheads indicate pale bodies. (**m**) Colocalization analysis of neuromelanin-positive neurons labeled with α-syn. (**n**) Colocalization analysis of neuromelanin-positive neurons labeled with P-α-syn. Results are mean ± SEM of N=3 IBD or control subjects and are expressed as percentage of dopaminergic neurons. Statistical analysis: Student-Newman-Keuls Method, with α=0.05, *p<0.01. (**o**) Representative immunohistochemical image (developed with DAB) from a patient of IBD, showing accumulation of α-syn (arrows). Scale bars: 30 µm (a-l); 40 (o)µm. Source data are provided as a Source Data file.

**Supplementary Figure 13.**
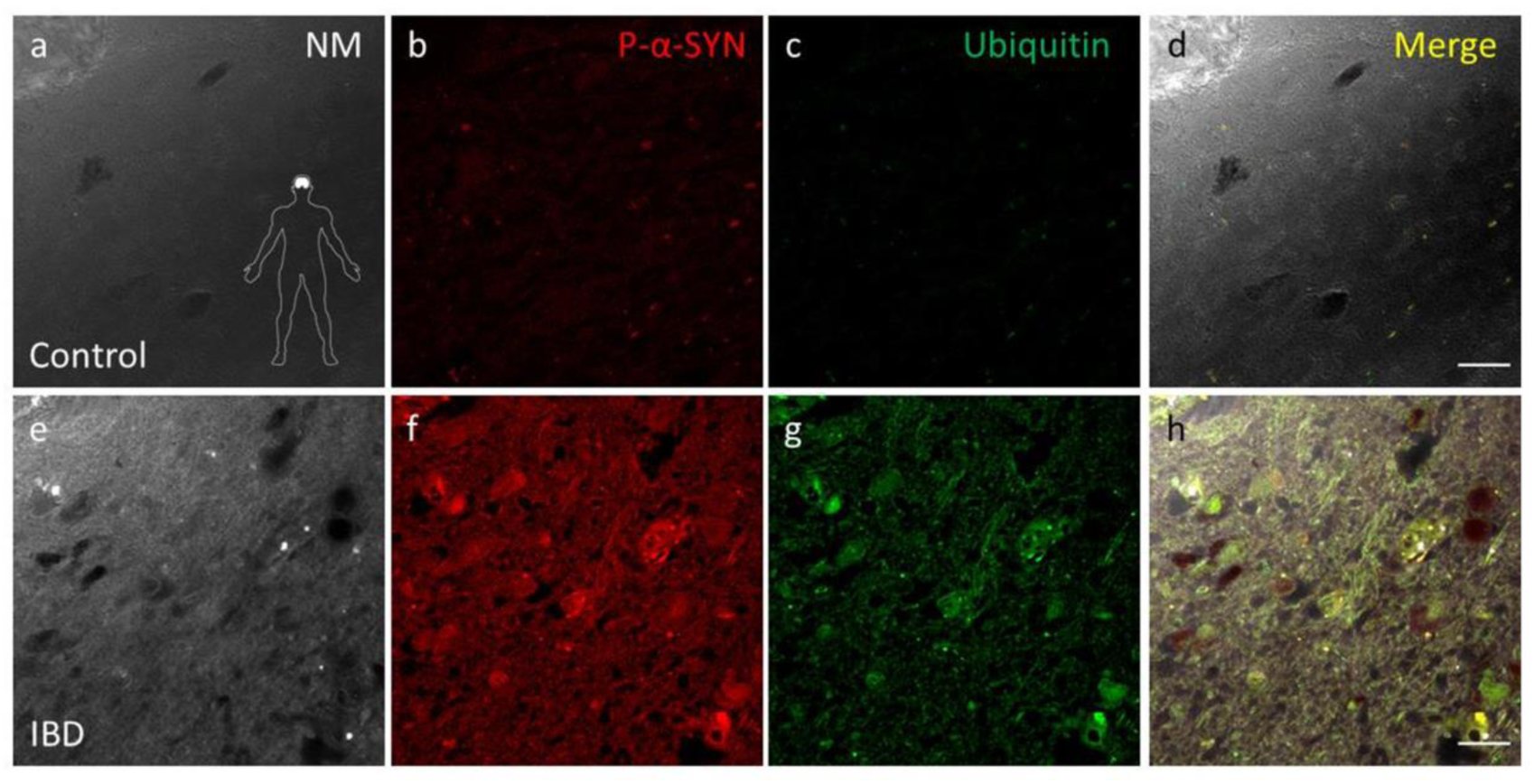
Immunofluorescence of P-α-syn and ubiquitin in mesencephalic neurons of control and IBD subjects. Microphotographs from the SN of control (**a-d**) and IBD subjects (**e-h**). **a** and **e** panels show bright fields with neuromelanin as dark structures; **b** and **f** panels show immunofluorescence to P-α-syn (red); **c** and **g** panels show immunofluorescence to ubiquitin (green); **d** and **h** panels correspond to merge of all images showed. Abbreviations: NM, neuromelanin; IBD, inflammatory bowel disease. Scale bars: 30 µm.

**Supplementary Figure 14.**
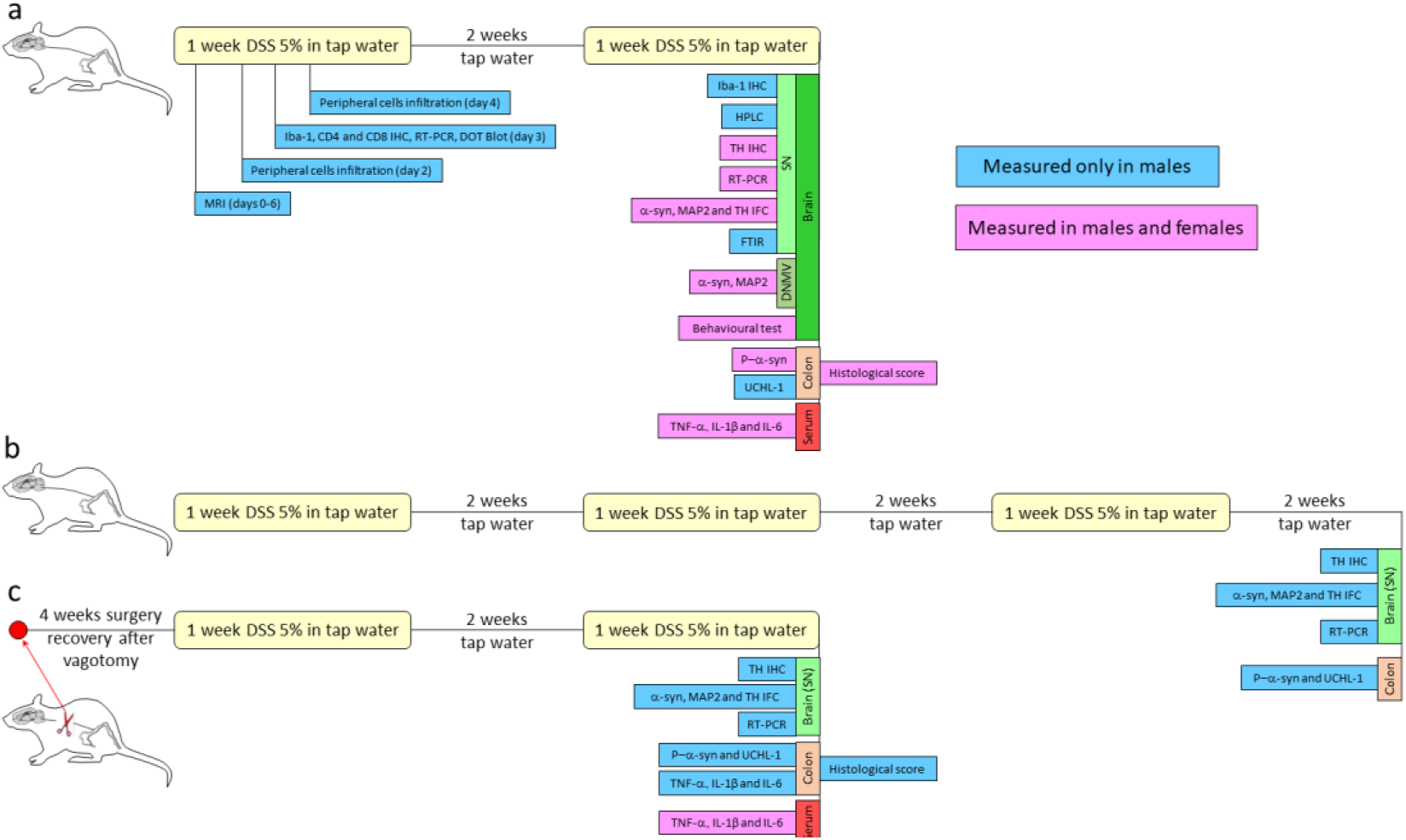
Timeline of the treatments with DSS in animals, and points at which the different parameters have been measured. (**a**) **Subchronic treatment with DSS.** Experimental animals (male and female Wistar rats) had free access to water with 5% DSS (yellow boxes) for 2 periods of a week separated by a period of two weeks in which they drank tap water. Control animals drank only tap water. At the end of the treatment, different parameters were measured to verify the effect of the subchronic treatment. Likewise, some parameters were measured at short times to determine the initial effects of the treatment with DSS. For the measurement of each parameter, control and experimental animals subjected during the same time to their respective treatments were used. **(b) Chronic treatment with DSS.** Experimental animals (male Wistar rats) had free access to water with 5% DSS (yellow boxes) for 3 periods of a week each followed by a period of two weeks in which they drank tap water. At the end of the treatment, different parameters were measured to verify the effect of the chronic treatment. **(c) Subchronic treatment with DSS plus vagotomy**. Male Wistar rats were vagotomized or sham-operated (red spot) and allowed 4 weeks to recover from surgery. Then they drank tap water for 4 weeks (controls) or underwent the 5% DSS paradigm described above. At the end of the treatment, different parameters were measured to check the effect of vagotomy on the treatment with DSS. The timeline of the control animals (consisting of 4 weeks drinking tap water) is not shown. Blue boxes represent the parameters measured only in males, whereas magenta boxes represent the parameters measured in both males and females.

**Supplementary Figure 15.**
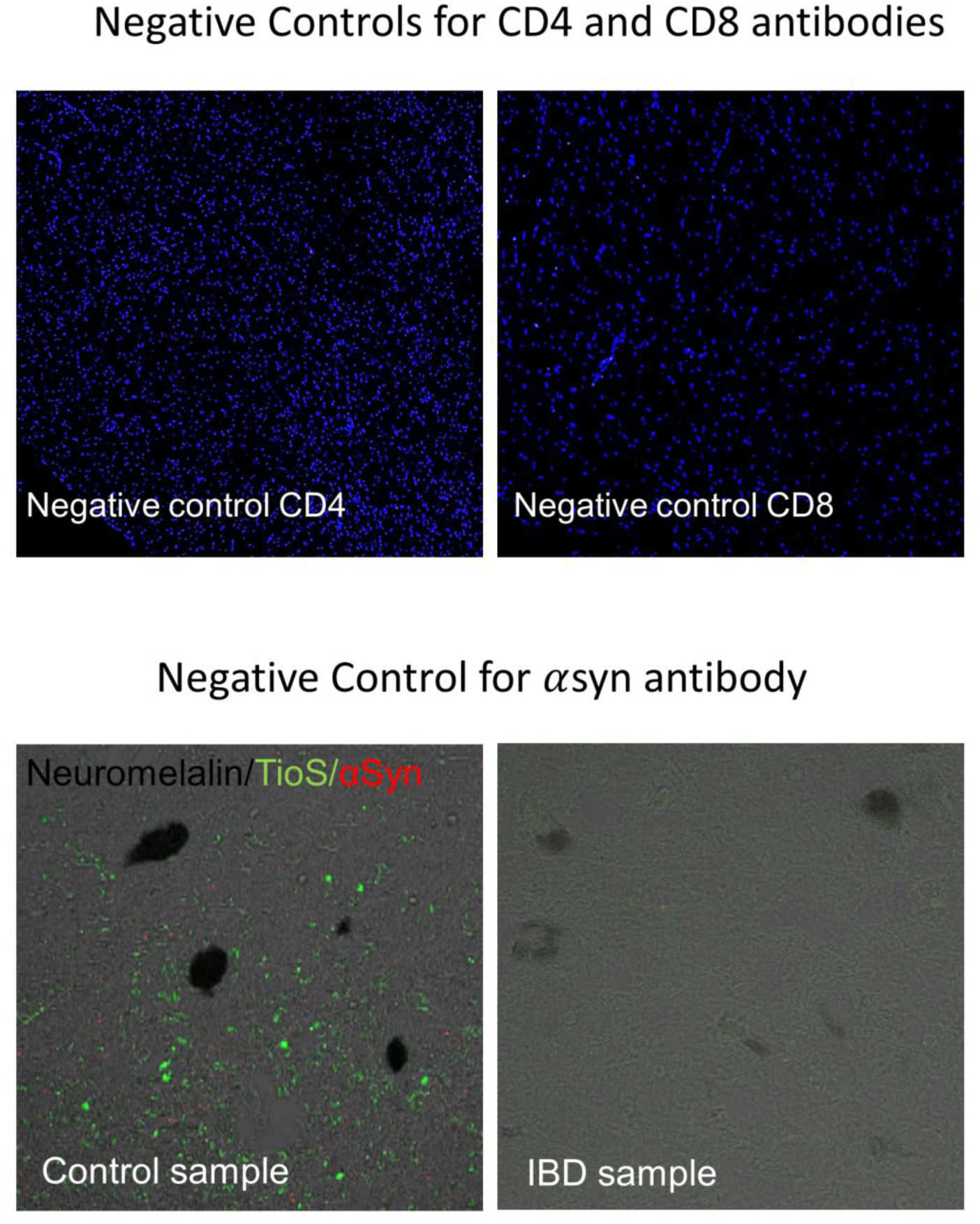
Negative controls for immunofluorescence. As a negative control, the primary step was omitted.

**Supplementary Table 1.** List of patients with ulcerative colitis and control patients used for gut studies.

|  | Case ID | Age | Sex | Diagnosis |
| --- | --- | --- | --- | --- |
| **Inflammatory bowel disease** | 13B0018342 | 33 | Male | Ulcerative colitis |
|  | 15B0003740 | 36 | Male | Ulcerative colitis |
|  | 13B0024824 | 46 | Female | Ulcerative colitis |
|  | 13B0031337 | 64 | Male | Ulcerative colitis |
|  | 16B0015532 | 65 | Female | Ulcerative colitis |
|  | 12B0012227 | 71 | Female | Ulcerative colitis |
| **Controls** | 15B0007953 | 31 | Female | Healthy colon |
|  | 15B0000707 | 49 | Male | Healthy colon |
|  | 15B0000054 | 60 | Female | Healthy colon |
|  | 14B0032988 | 65 | Male | Healthy colon |
|  | 15B0000767 | 74 | Female | Healthy colon |
|  | 15B0004710 | 83 | Male | Healthy colon |

**Supplementary Table 2.** List of patients with inflammatory bowel disease and control patients used for brain studies.

|  | Case ID | Age | Sex | Diagnosis |
| --- | --- | --- | --- | --- |
| **Inflammatory bowel disease** | 05/018 | 53 | Male | Ulcerative colitis. Multiple Sclerosis. Sepsis due to Gram-negative organism. |
|  | 03/102 | 45 | Male | Ulcerative colitis. Circumscribed brain atrophy |
|  | 09/086 | 38 | Female | Gastroenteritis and Colitis. Spinal muscular atrophy |
|  | 99/2009 | 59 | Male | Crohn's Disease. Lung Cancer |
|  | B1737 | 52 | Female | Ulcerative Colitis. Dystrophia myotonica |
|  | A11-33 | 37 | Male | Crohn's Disease |
|  | 20.027-1 | 65 | Male | Crohn's Disease. Septic shock |
|  | 20.027-2 | 40 | Female | Ulcerative Colitis. Cerebral hemorrhage |
| **Controls** | 13/037 | 51 | Male | Chronic Obstructive Pulmonary Disease |
|  | 13/039 | 41 | Male | Normal Brain |
|  | 13/064 | 57 | Female | Lung Cancer |
|  | 13/103 | 48 | Female | Normal Brain |
|  | 13/156 | 45 | Male | Myocardial infarct |
|  | A11-2 | 35 | Male | Acute pneumonia |
|  | 20.027-5 | 54 | Male | Multiple myeloma |
|  | 20.027-7 | 36 | Male | Septic shock |

**Supplementary Table 3.**
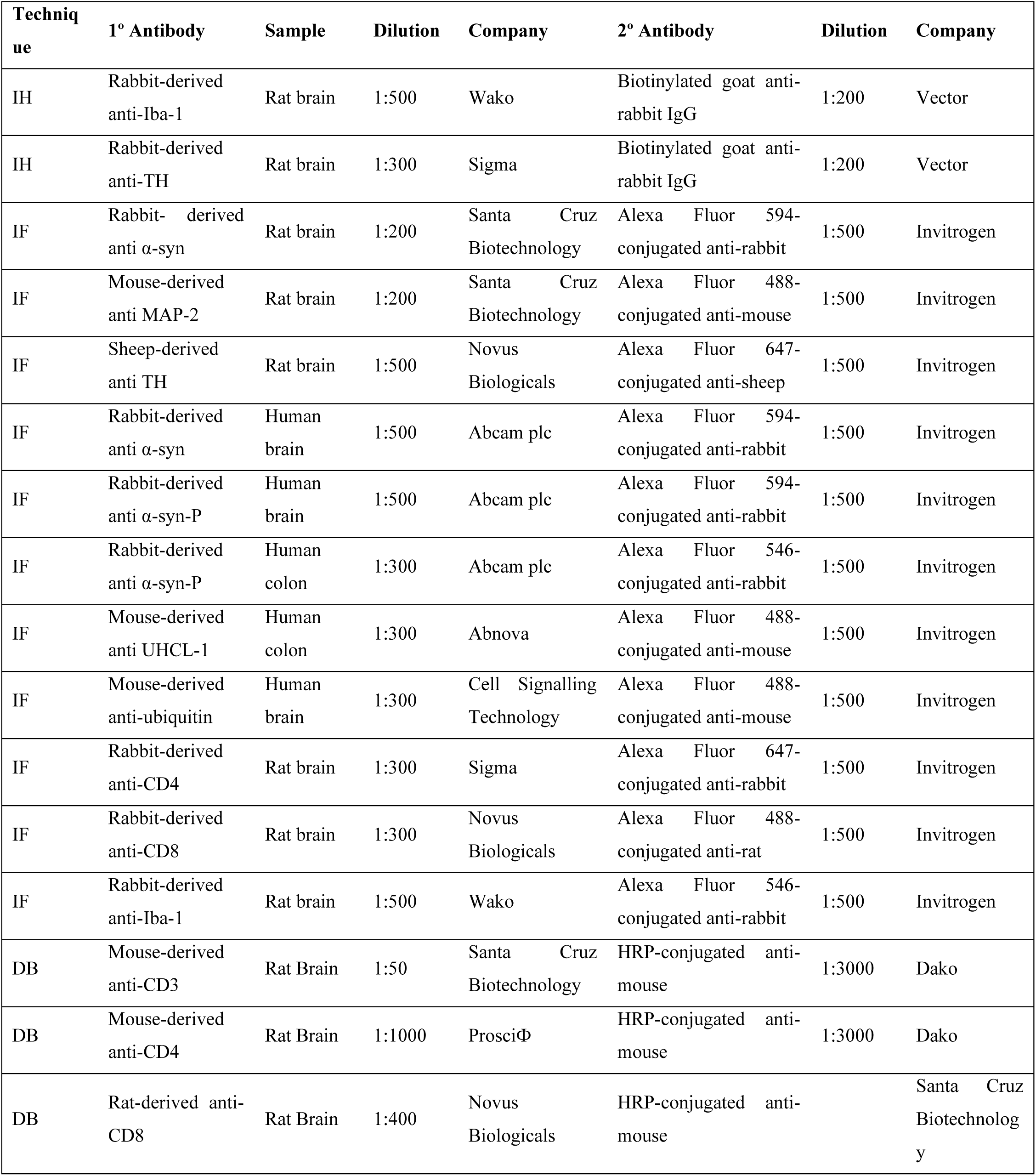
Primary and secondary antibodies used for immunohistochemistry (IH) and immunofluorescence (IF).

**Supplementary Table 4.** Antibody panels used for flow cytometry

| **Tube** | *FITC* | *PE* | *PE Cy7* | *APC* |
| --- | --- | --- | --- | --- |
| **1** | CD45 | CD4 | CD8 | CD3 |
| **2** | CD45 | CD11b | CD8 | CD3 |

**Supplementary Table 5.**
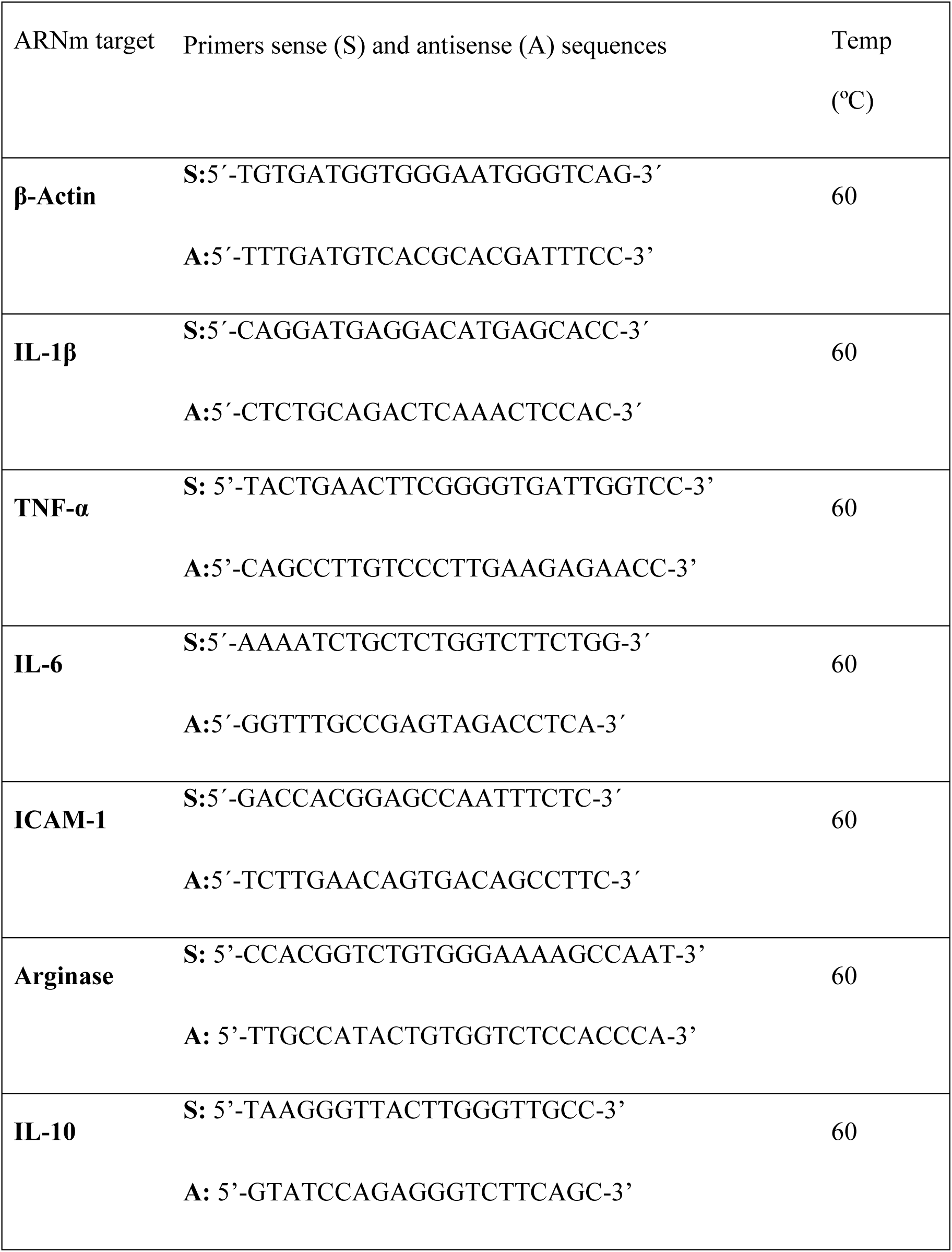
Primers used for real-time PCR.

